# Dynamic nucleosome redistribution and increases in nucleosome sensitivity underpin THP-1 macrophage response to LPS

**DOI:** 10.1101/2025.02.11.637695

**Authors:** Jane M Benoit, Brandon D Buck, Mahdi Khadem, Hank W Bass, Jonathan H Dennis

## Abstract

Macrophages detect lipopolysaccharide (LPS) through toll-like receptor 4 (TLR-4) on the cell surface which initiates a signaling cascade resulting in the recruitment of regulatory factors to chromatin and subsequent expression of chemokine and cytokine genes. Primary response genes, marked by poised promoters and enhancers, are rapidly expressed after LPS stimulation, and their gene products activate secondary response genes via paracrine and autocrine signaling pathways. While the signaling cascades following macrophage activation are well understood, the dynamics of nucleosome architecture and regulatory factor binding in promoter regions during early and late LPS responses remain unclear. Here, we stimulated THP-1 derived macrophages with LPS and assessed nucleosome distribution and MNase sensitivity across promoters at eight time points spanning primary and secondary responses. We found that while nucleosome distribution was static over most promoters, LPS stimulation resulted in transient remodeling of a subset of immune and DNA repair gene promoters. We also observed distinct MNase sensitivity alterations in two phases which aligned with early and late gene expression patterns. Notably, while most Pol II promoters showed altered chromatin sensitivity, only a subset exhibited transcriptional changes, suggesting that widespread alterations in nucleosome distribution and sensitivity occur at promoters with or without alterations in gene expression. These findings provide new temporal insights into the transient and long-term effects of immune stimulation on promoter architecture and offer a methodological framework for additional time-resolved studies of chromatin remodeling in other systems.

**Summary sentence:** Following LPS stimulation, a subset of nucleosomes in macrophage immune promoters undergo transient redistribution, whereas the majority of nucleosomes show changes in MNase sensitivity that are largely uncoupled from gene expression.

Graphical abstract.
LPS stimulation of THP-1 derived macrophages leads to altered nucleosome occupancy and positioning within a subset of promoters which are enriched for LPS response genes. Altered distribution patterns permit regulatory factor binding and gene expression. The majority of promoters have altered nucleosome sensitivity with a trend towards increased sensitivity after LPS stimulation. Altered sensitivity results in regulatory factor binding and expression of LPS response genes.

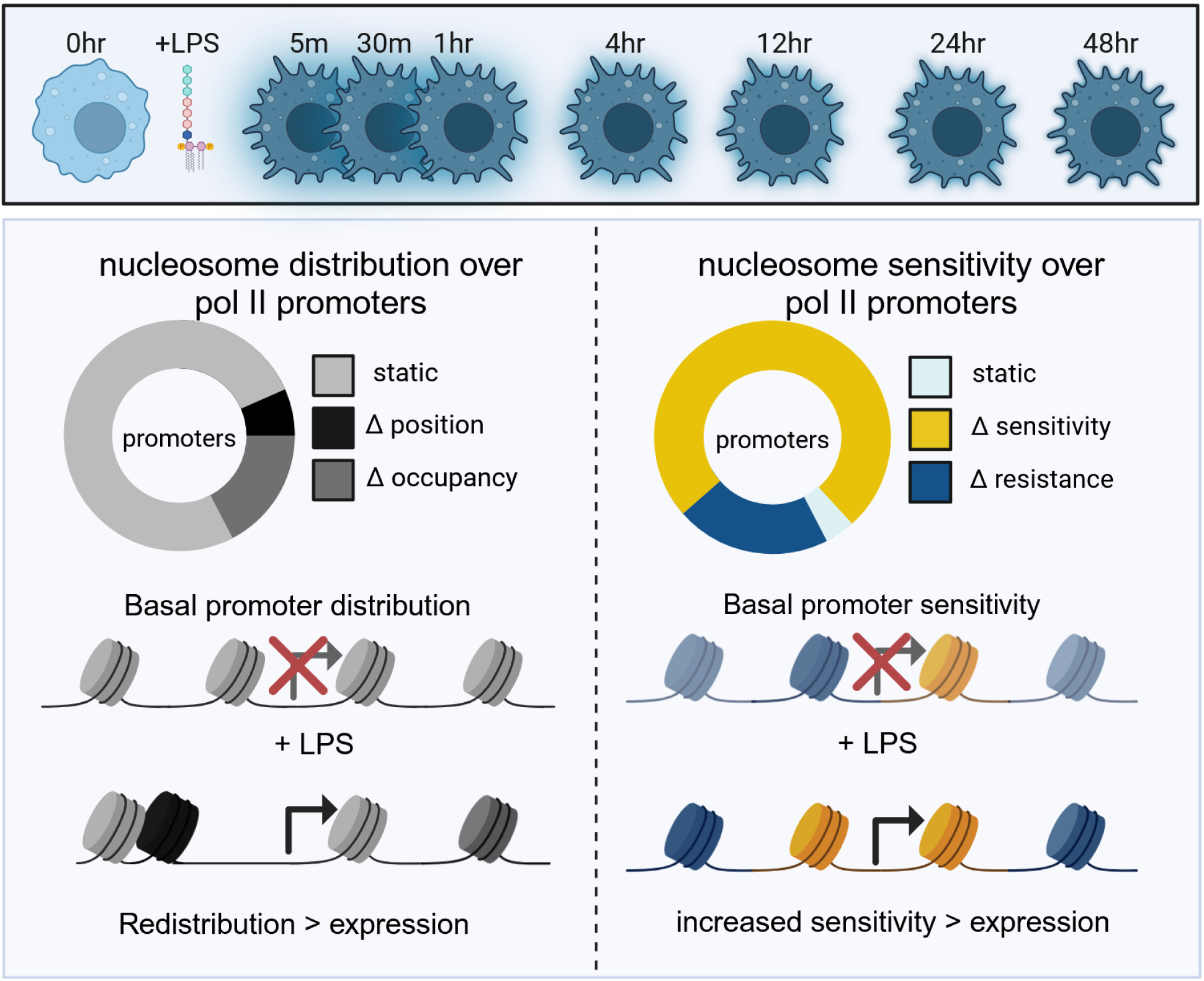

## Introduction

Macrophages are key innate immune cells which are primed to recognize and respond to pathogens in a rapid and highly specific manner. Macrophages are derived from a common myeloid progenitor and are present in almost all tissue types ^1,2^. These cells express pathogen recognition receptors (PRRs), including Toll-like receptors (TLRs), which recognize microbial signatures and initiate signaling cascades leading to rapid cellular responses^3–7^. TLR mediated ligand detection activates signaling pathways mediated by combinations of intermediate proteins, such as MyD88 and TRIF, which leads to downstream signaling and translocation of nuclear factor κB (NF-κB) and/or interferon regulatory factors (IRFs) to the nucleus where they activate expression of type I interferon, chemokine, and cytokine genes ^8–11^. TLR-4, which recognizes lipopolysaccharide (LPS) on the surface of gram negative bacteria and selected damage associated ligands, is highly expressed by macrophages and other myeloid cells ^12,13^. TLR4, dimerized with MD-2, binds LPS to initiate intracellular signaling through MyD88 and TRIF which results in AP-1, NF-κB, and IRF mediated gene activation ^10,14–16^. In addition to immediate ligand detection and signaling, macrophages can become polarized or tolerized by immune stimulation which results in more nuanced tuning of immune responses to future stimuli^5,17.^

TLR signaling results in rapid translocation of pre-formed regulatory factors from the cytoplasm to the nucleus where they target specific binding motifs at immune responsive genes. Primary response genes are activated by the initial influx of regulatory factors within one hour of induction and products of these genes stimulate secondary response genes are activated several hours later due to the need for chromatin remodeling and protein synthesis ^18–20^. The timing and scope of the macrophage immune response is mediated by chromatin structure with primary response gene products participating in autocrine and paracrine signaling pathways ^21–28^. During macrophage development, chromatin is restructured over the enhancers and promoters of primary response genes with alterations in histone post translational modifications (HPTMs), chromatin looping, DNA methylation, chromatin accessibility, and nucleosome distribution which posies these genes to respond upon TLR stimulation ^21–26^. These alterations in chromatin structure physically open transcription factor (TF) binding sites to allow motif recognition and binding which permits induction of transcription ^5,26,28–30^. The enhancers of primary response genes are poised to respond to immune stimulation in mature macrophages as defined by accessibility and histone modification profiles associated with poised but inactive states ^30–33^. The promoters of these genes are similarity primed to respond to stimulation with a combination of active marks, accessible regions, unstable nucleosomes over CpG islands, and paused polymerases ^18,20,34,35^.

While primary response genes are poised prior to stimulation, secondary response genes must undergo extensive chromatin remodeling prior to expression after TLR-4 stimulation. The BRG1/BRM subunits of the SWI/SNF class of ATP-dependent nucleosome remodeling complexes are required for activation of many secondary response genes ^19^. Hundreds of latent enhancers are inactive in unstimulated macrophages and only acquire TFs and appropriate HPTMs after immune stimulation ^36,37^. Additionally, alterations in higher order chromatin structure and dynamic looping are required for appropriate induction of innate immune responses ^33,38,39^. Local alterations in nucleosome positioning, HPTM localization, and chromatin accessibility have been observed in both enhancers and promoters following TLR-4 stimulation ^20,34,40–43^. In particular, binding of NF-κB and IRFs have been strongly associated with chromatin remodeling and subsequent expression ^35,40,42^.

Nucleosomes, consisting of ∼146 bp of DNA wrapped around a core histone octamer, are the fundamental subunit of chromatin and can be shifted or evicted from chromatin to allow for regulatory factor binding and recruitment of transcriptional machinery ^29,44,45^. The importance of nucleosomes within promoters is illustrated by the ability of nucleosomes to prevent the initiation of transcription by preventing the pre-initiation complex assembly ^46^ and that depletion of nucleosomes results in expression of otherwise silent genes^47^. Promoter nucleosomes can be fully displaced by the passage of Pol II ^46,48,49^ and active promoters are marked by a pronounced nucleosome depleted region which is permissive to transcriptional machinery and a positioned +1 nucleosome ^50–52^. The location of nucleosomes across promoters can be mapped using Micrococcal nuclease (MNase) digestion of crosslinked chromatin ^53–57^. Additionally, titration of MNase digestion has resulted in the identification of nucleosomes which are sensitive or resistant to digestion ^58–63^. This measure of MNase sensitivity corresponds with expression levels and transcription factor binding ^60–62,64,65^ suggesting that it is an important feature that can distinguish active and inactive regions of the genome.

While much is known about the innate immune response to TLR-4 stimulation, there are still open questions about the dynamics of nucleosome remodeling and how specific regulatory factors interact with proinflammatory genes. Namely, how rapidly, and to what extent, do promoters respond to TLR-4 induction and what is the extent and kinetics of promoter recovery after stimulation. While previous studies have focused on primary and secondary responses, few time course experiments capture the very earliest stages of chromatin remodeling over promoters or stretch into the days following immune stimulation ^18–20,33,35,40,42^. Additionally, many studies have focused on alterations in chromatin accessibility, as measured by ATACseq, as a proxy for chromatin remodeling without assaying underlying nucleosome architecture ^33,35^. While insightful for TF binding, this method does not identify the location of underlying nucleosomes and the measured accessibility may result from a combination of chromatin features rather than a discrete remodeling event ^42,66,67^. The time-resolved chromatin remodeling data resulting from TLR-4 immune stimulation in macrophages provides insight into the ordered events leading to nucleosome displacement, regulatory factor binding, gene expression, and recovery in a highly consistent model.

We used the well established THP-1 macrophage model ^68^ to examine the role of nucleosome and regulatory factor dynamics during early and late immune responses to LPS. Despite being an immortalized cell line, THP-1 derived macrophages express near normal levels of immune signaling receptors and stimulation with LPS results in strong cytokine production and phagocytic activity ^69,70^ making them a useful tool for understanding macrophage responses to immune stimuli. We treated THP-1 derived macrophages with 10ng/ml LPS and assayed nucleosome distribution and nucleosome sensitivity to MNase digestion at eight time points spanning the early and late stages of LPS stimulation. We found that nucleosome positioning is largely stable over most pol II promoters with transient alterations in nucleosome distribution at a subset of immune response genes after LPS stimulation. In contrast, nucleosome sensitivity to MNase digestion was highly dynamic and did not return to a basal state within 48 hours after stimulation. This result suggests that while alterations in nucleosome distribution are largely localized and transient, alterations in nucleosome sensitivity are more durable and may underpin future polarization or tolerization states. Together, these findings demonstrate that TLR-4 stimulation produces rapid and transient alterations in nucleosome distribution and lasting alterations in sensitivity to MNase digestion corresponding to primary and secondary immune responses.

## Methods

### Tissue culture and nuclei isolation

THP-1 monocytes were obtained from the ATCC (TIB-202) and were cultured in RPMI-1640 media (Gibco) supplemented with 10% fetal bovine serum and 1% penicillin-streptomycin. Cells were maintained at 37°C with 5% CO2. Cells were exposed to 25nM phorbol 12-myristate 13-acetate (Sigma-Aldrich) for 48 hours and recovered in complete medium for 24 hours following differentiation ^69^. Differentiated cells were treated with 100 ng LPS and harvested zero minutes, five minutes, 30 minutes, one hour, four hours, 12 hours, 24 hours, and 48 hours after stimulation. At each time point, cells were crosslinked in 1% v/v formaldehyde (Sigma-Aldrich) and incubated for 10 minutes at room temperature while rocking. Excess formaldehyde was quenched with 125mM glycine and cells were washed twice in PBS. Cells were resuspended in nuclei isolation buffer (10 mM HEPES at pH 7.8, 2 mM MgOAc^2^, 0.3 M sucrose, 1 mM CaCl^2^ and 1% Triton-X) and pelleted by centrifugation at 1000 × *g* for 10 min at 4 °C. Nuclei were washed twice in nuclei isolation buffer and nuclei purity was confirmed by DAPI staining.

### MNase cleavage and preparation of DNA libraries

All nuclei samples were processed in replicate. For each reaction, 2.5x10^6^ nuclei were resuspended in 500ul nuclei isolation buffer (10 mM HEPES at pH 7.8, 2 mM MgOAc^2^, 0.3 M sucrose, 1 mM CaCl^2^ and 1% Triton-X) and digested with Micrococcal nuclease (MNase) (Worthington) under light (5U) or or heavy (200U) conditions for 10 minutes at 37 °C. MNase reactions were halted with 50 mM EDTA. Protein-DNA crosslinks were reversed by treating the MNase-digested nuclei with 0.2 mg/mL proteinase K and 1% sodium dodecyl sulfate, and incubating overnight at 65 °C. DNA was isolated using guanidinium-isopropanol and washed with ethanol on a silica column. Mononucleosomal and sub-mononucleosomal DNA was isolated from a 2% agarose gel slice by excising all fragments under 200bp in length and purified by glycogen precipitation.

MNase sequencing libraries were prepared for each replicate using the Accel-NGS®2S Plus DNA Library Kit (Swift) for Illumina® using 10ng of input DNA from each digestion. Libraries were indexed with the Accel-NGS 2S Unique Dual Indexing Kit and normalized with Normalase. Library size and quality was verified with a D1000 Tapestation. Molar concentration of each indexed library was determined by KAPA quantitative PCR and size corrected using sizing information from the Tapestation.

### Solution-based sequence capture and Illumina sequencing

We used a previously validated ^54,56^ custom designed Roche Nimblegen SeqCap EZ Library SR to capture 2000 bp regions flanking the TSS for every gene in the human genome. The TSS sequences were repeatedly masked using human COT DNA and library adapters were masked with universal blocking oligos. Biotinylated sequence capture oligos were hybridized to input libraries for 48 hours and purified with streptavidin beads. Enriched sequences were amplified in a 15 cycle PCR amplification using TruSeq primers. Final library size and quality was validated with the Tapestation and KAPA quantitative PCR was used to determine final molar concentrations. Paired end 50 bp sequences were generated on the FSU College of Medicine’s NovaSeq 600 Illumina sequencer.

### Nucleosome data processing

Raw sequencing reads were demultiplexed using Cassava and adapter sequences and poor quality reads were removed with Trimmomatic (version 0.39)^71^. Bowtie2 was used to align the sequences to the T2T reference human genome with the --end-to-end --no-mixed --no-discordant flags^72^. Mitochondrial and duplicate reads were filtered with samtools (version 1.10) and picardtools (version 2.27.3)^73^. Nucleosome footprints for light and heavy digests were calculated with deeptools (version 3.5.1) and DANPOS (version 3.1.1)^74,75^. The log2 ratio of light to heavy digests was calculated for each time point to determine nucleosome sensitivity to MNase digest. Figures showing individual loci were generated with the PlotGardener package (version 1.4.1) in R (version 4.3.2) and the UCSC genome browser 76^76,77^. Additional processing of bigwig files to generate z-score matrices and upset plots was done in R (version 4.3.2), bash, and python3 with custom scripts. Individual genes were identified and grouped by gene ontology using Enrichr to better understand changes across related loci^78^. Nucleosome occupancy and sensitivity files were uploaded to the UCSC genome browser and the NCBI GEO database under accession number GSE279622.

### Public data processing

External data from the GEO dataset GSE147314^35^ and GSE201376 ^33^ were obtained from NCBI and concurrently processed. RNAseq data was trimmed with Trimmomatic^71^, aligned to the T2T reference genome with HiSat2^79^, and further processed with Stringtie^80^ and Ballgown to call differentially expressed genes. ATACseq and ChIPseq data were trimmed with trimmomatic^71^ and aligned to the T2T reference genome with Bowtie2^72^. Mitochondrial and low quality reads were filtered with samtools^73^ and duplicate reads were removed with picardtools and peaks were called with MACS3^81^.

### Tipps, Tricks, and Traps

The level of digestion used in MNase based assays is critical for classification of sensitive and resistant nucleosomes. Our digestions were optimized for 2.5x10^6^ THP-1 cells per reaction. Digestion time, reaction temperature, amount of enzyme, cell count, and total volume will impact digestion levels and may need to be optimized to different sample types. We aim for a full nucleosomal ladder extending from mononucleosomes to large fragments at the limit of resolution for our light digests. This is achieved by a titrating MNase until the lightest possible digest is achieved that meets these conditions. For heavy digests, we aim for a nucleosomal ladder that shows mononucleosomes and up to penta and hexa nucleosome arrays. A heavy digest should not have any large fragments at the limit of resolution on an agarose gel. We recommend testing digest conditions on untreated cells to optimize conditions for different cell types and experimental conditions.

## Results

### LPS stimulation results in nucleosome re-distribution over a subset of pol II promoters

To understand promoter architecture during early and late innate immune responses, we stimulated THP-1 derived macrophages with lipopolysaccharide (LPS). We then assayed promoter nucleosome distribution and sensitivity to MNase digestion (summarized in Fig. 1). Samples were collected at eight timepoints: 0 hour, 5 minutes, 30 minutes, 1 hour, 4 hours, 12 hours, 24 hours, and 48 hours post-stimulation. These timepoints were selected to capture periods of chromatin remodeling associated with early and late immune responses ^18–20,82^.

**Figure 1.**
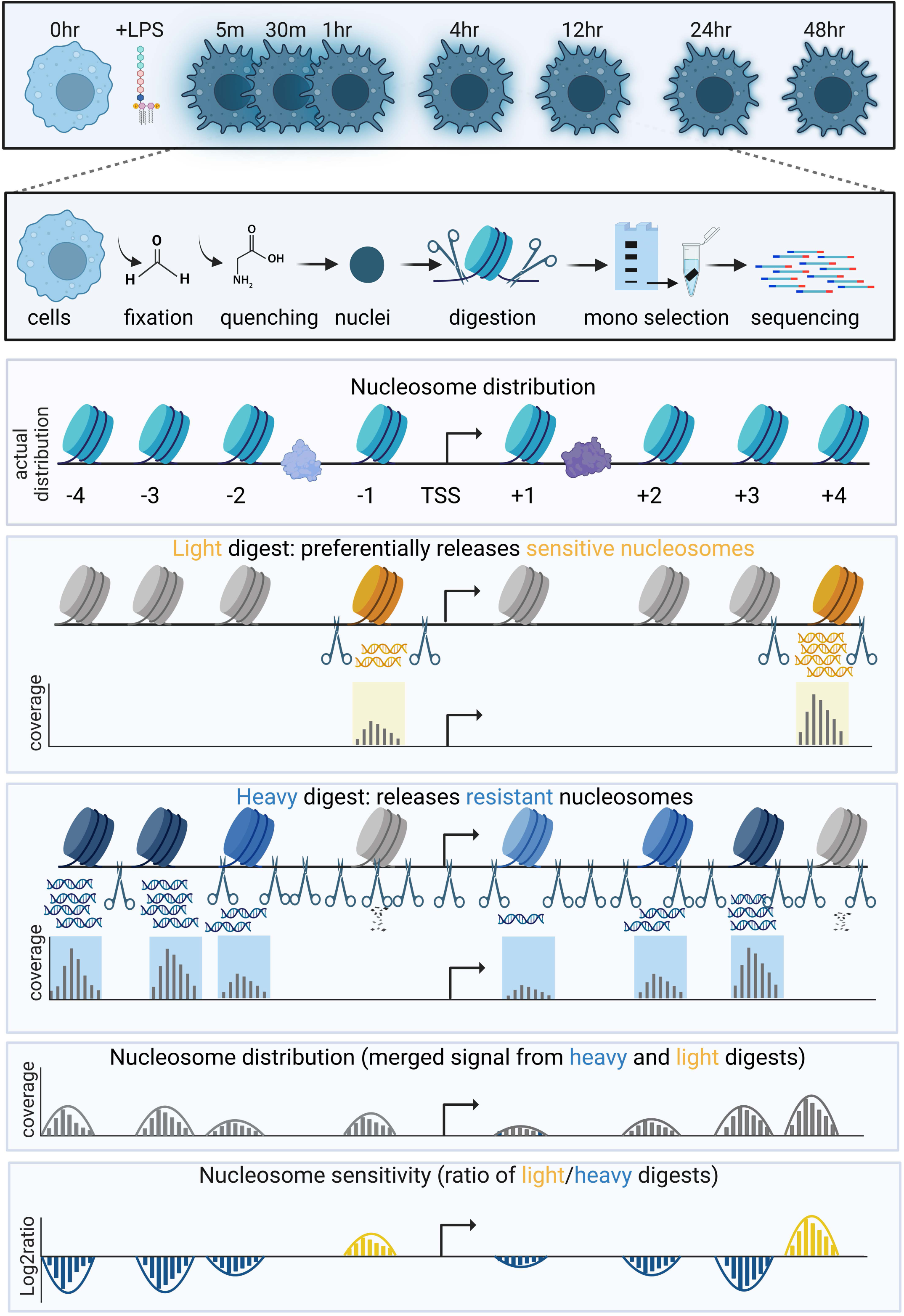
Schematic of MNased assays used to profile macrophage chromatin after LPS stimulation. A. THP-1 derived macrophages were sampled before and 5min, 30min, 1hr, 4hr, 12hr, 24hr, and 48hr after treatment with LPS. Cells were fixed in formaldehyde which was quenched with glycine. Isolated nuclei were digested with MNase and mononucleosomal DNA was sequenced. Light MNase digestion releases a subset of sensitive nucleosomes (gold) while heavy MNase digestion preferentially releases resistant nucleosomes (dark blue). Heavy conditions can over digest sensitive nucleosomes (shown as shredded nucleotides). Nucleosome distribution is determined by combining mononucleosomal reads between 120-180 bp from light and heavy digests. Nucleosome sensitivity is determined by taking the log2 ratio of light/heavy digests. Sensitive nucleosomes (yellow) are overrepresented in light digests while resistant nucleosomes (blue) are overrepresented in heavy digests.

We used high and low concentrations of MNase to produce heavy and light digests (Fig. 2A), allowing us to profile both nucleosome distribution and sensitivity to MNase digestion (Fig. 1). We combined the 120-180bp fragments from both digests to profile overall nucleosome distribution, irrespective of digestion levels ^58,62,63,83^. The ratio of fragments from light and heavy digests was used to determine nucleosome sensitivity to MNase digestion (described in ^58^).

**Figure 2.**
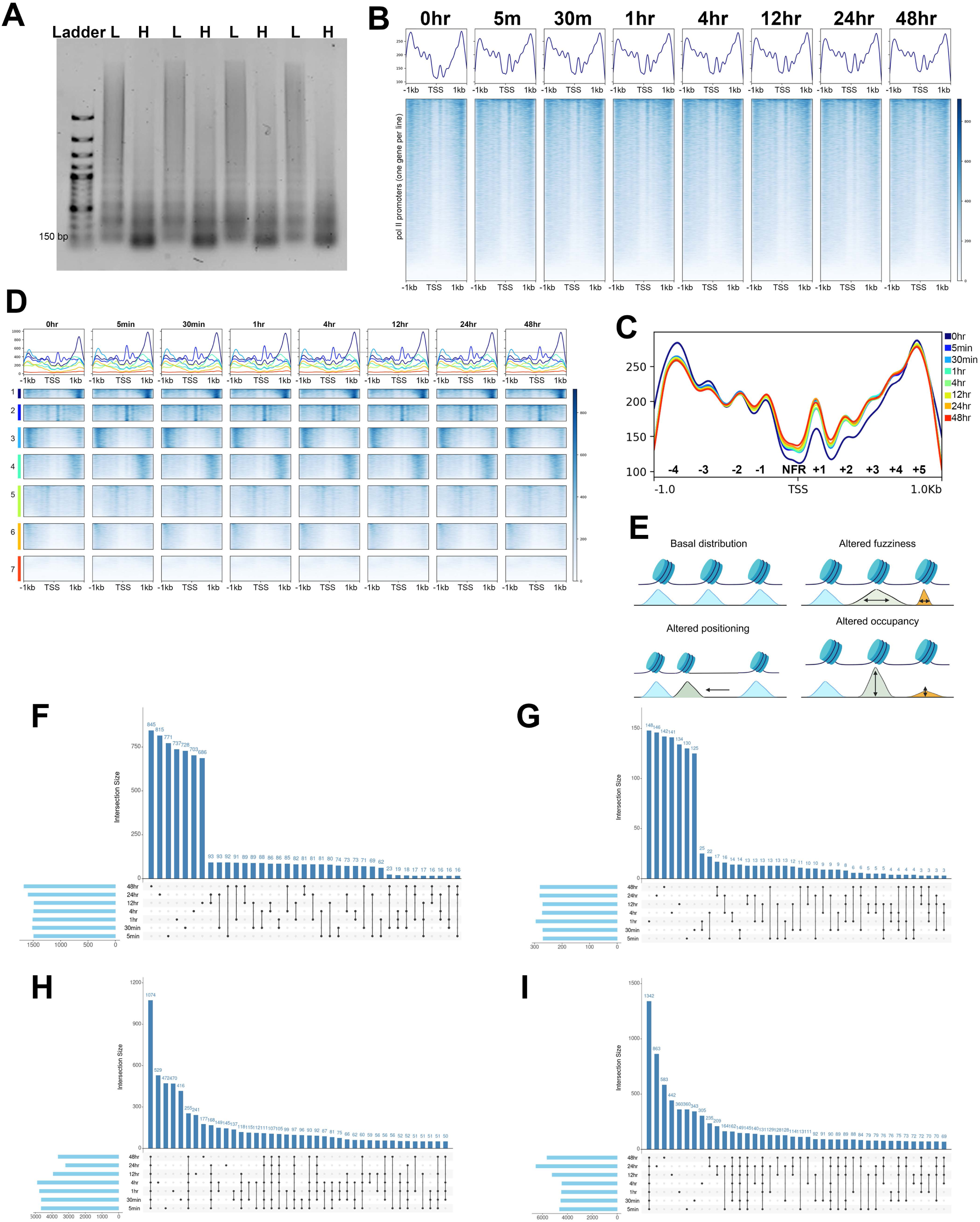
Nucleosome distribution over a select set of pol II promoters is impacted by LPS stimulation. A. Representative agarose gel showing DNA isolated from heavy and light MNase digests of crosslinked chromatin. L indicates light digests and H indicates heavy digests. B. Heatmaps show nucleosome distribution signal in blue over a 2kb region centered on the TSS of 21,054 pol II genes. Line plots above each map show average signal values. C. Average nucleosome distribution patterns across all time points overlaid to show defining promoter features. 95% confidence intervals are shown as shadows behind each average line. D. Nucleosome distribution profiles for all time points clustered into seven groups by kmeans clustering based on the 0hr time point. The dashed line indicates the signal of the +1 nucleosome signal in unstimulated macrophages. E. Summary of nucleosome distribution alterations. Basal positions and occupancy are shown in pale blue with alterations in fuzziness, positioning, and occupancy indicated by arrows and color changes. F. Upset plot showing common and unique genes with repositioned nucleosomes. G. Upset plot showing common and unique genes with altered nucleosome fuzziness profiles. H. Upset plot showing common and unique genes with increased nucleosome occupancy. I. Upset plot showing common and unique genes with decreased nucleosome occupancy.

We conducted high resolution mapping of MNase protected nucleosomes in a 2kb window centered on the transcription start sites (TSSs) of pol II RefSeq promoters using the previously described mTSS-seq method ^54,56^. This method provides high coverage of promoter regions with as few as 20 million reads per sample. To verify the effectiveness of our mTSS-seq approach, we mapped nucleosome occupancy over a 4kb region spanning the promoters of pol II genes. The results showed significant enrichment over the immediate promoter region (Supp. Fig. 1A), with greater coverage over the 2kb region centered on the TSS compared to the flanking regions.

We were first interested in determining if nucleosome distribution over promoters was altered by LPS stimulation. Accordingly, we plotted nucleosome distribution over pol II promoters (Fig. 2B) and found that the average distribution profile was highly similar at all time points. We performed a correlation analysis of nucleosome distribution in 10bp bins across all promoters. The Pearson’s correlation coefficient was greater than 0.87 for all samples (Supp. Fig. 1B), indicating highly similar profiles for most promoters, with unstimulated macrophages as the outlier. Direct comparison of average profiles revealed greater overall nucleosome occupancy downstream of the TSS following LPS stimulation, with a significant increase in occupancy over the +1 nucleosome (Fig. 2C). As these profiles were generated from aggregated promoter data, we were interested if groups of promoters had similar architectural features. We used k-means clustering to classify promoters by dominant nucleosome distribution patterns within 2kb of the transcription start sites (Fig. 2D). Similar promoter classes were observed at all time points, with a notable increase in occupancy over the +1 and +2 nucleosomes following LPS stimulation, persisting through 48 hours (Fig. 2D, most apparent in average plots). This suggests that while the majority of nucleosomes in pol II promoters have static distribution patterns, a subset of nucleosomes have altered profiles following LPS stimulation.

We wanted to identify and characterize the nucleosome landscape of promoters with altered nucleosome distribution following LPS treatment. Our previous work with reactivation of Kaposi’s Sarcoma Associated Herpesvirus (KSHV) showed that a subset of promoters undergo rapid and transient alterations in nucleosome positioning ^54,55^. To determine if the same phenomenon applied to LPS stimulation of macrophages, we used DANPOS3^74^ to identify promoters with altered nucleosome distribution, defined by position, fuzziness, and occupancy.

Our first measure of nucleosome distribution, nucleosome positioning refers to the precise location of a nucleosome in relation to underlying DNA (Fig. 2E). We defined repositioning as a shift in the nucleosome summit position greater than 80bp between time points. We compared the positions of all promoter nucleosomes at each time point after LPS stimulation with the basal position in unstimulated cells. We identified over 1,400 genes with one or more nucleosomes within promoters which were repositioned following LPS treatment at any given time point compared to basal positions (Fig. 2F, Table 1). Approximately 80% of these repositioning events were unique to a single time point or shared between three or fewer time points indicating that most repositioning events are transient and temporally regulated (Fig. 2F, Supp. Table 1). Interestingly, there was a trend of increasing repositioning events in the 24hr and 48hr time points compared to earlier time points (Table 1). This indicates two phases of reorganization of promoters as cells initially respond to the stimulus and later recover or adopt a more polarized state 24 hours after initial stimulation. To determine the functional relevance of repositioned nucleosomes, we used the Enrichr gene ontology tool^78^ to identify significantly enriched terms associated with promoters showing altered nucleosome positioning after LPS stimulation (Supp. Fig. 2A). Repositioned nucleosomes within 5 minutes of LPS stimulation were enriched for ephrin signaling which involves receptor protein-tyrosine kinases on macrophage surfaces and modulate TLR responses in dendritic cells ^84,85^. Genes associated with TLR signaling were enriched 30 minutes, 4hr, and 12hr after stimulation (Supp. Fig. 2A). Interestingly, genes involved in IL-27 signaling were selectively enriched 48hr after stimulation suggesting a trend towards pro-inflammatory polarization ^86–88^. While not significantly enriched, genes in the IL-2, IL-5, AP-1, and NF-KB signaling pathways were also identified as having repositioning events following LPS stimulation (Supp. Fig. 2A). The enrichment of immune pathways in these timepoints confirms that remodeling is targeted to a subset of relevant promoters following LPS stimulation.

**Table 1.** Number of genes with one or more re-distributed nucleosomes within 2kb promoter regions following LPS stimulation.

| Time | repositioning | altered fuzziness | increased occupancy | decreased occupancy |
| --- | --- | --- | --- | --- |
| 5min | 1489 | 269 | 4649 | 4633 |
| 30min | 1512 | 270 | 4641 | 4533 |
| 1hr | 1507 | 295 | 4752 | 4476 |
| 4hr | 1489 | 272 | 4877 | 4443 |
| 12hr | 1484 | 270 | 3928 | 5227 |
| 24hr | 1592 | 281 | 3184 | 6529 |
| 48hr | 1668 | 280 | 3630 | 5642 |

Our second measure of nucleosome distribution, nucleosome fuzziness refers to the broadness of the nucleosome peak irrespective of the summit height or the position of the nucleosome (Fig. 2E). Nucleosomes with lower fuzziness scores have a tighter based and more defined peak while those with higher scores have a less defined peak. We considered a change in nucleosome fuzziness to occur if there was a log2 fold change greater than 1.5 between conditions combined with a p-value and FDR under 0.05. We identified ∼300 nucleosomes with significantly altered fuzziness values at any given time point following LPS stimulation (Fig. 2G, Table 1). Similar to repositioning, over 70% of promoters with altered fuzziness were found at fewer than three time points following stimulation (Fig. 2G, Supp. Table 2) indicating that altered fuzziness occurs transiently. We found that genes with fuzziness alterations within five minutes of LPS stimulation were enriched for integrin interactions which are important for macrophage infiltration and polarization ^89,90^ (Supp. Fig. 2B). Promoters with altered fuzziness were enriched for a broad array of signaling pathways at 30 minutes including syndecan-1, Class I PI3K, LPA4, Hepatocyte Growth factor, Angiopoietin receptor, and androgen receptor pathways (Supp. Fig. 2B). Genes associated with HDAC mediated signaling events were enriched at 1hr and 4hr post stimulation while IL-12, JNK, IFN-γ, IL-6, and IL-2 pathways were enriched 24hr after LPS stimulation (Supp. Fig. 2B). This suggests that fundamentally different classes of genes experience altered nucleosome fuzziness in early and late response to LPS stimulation.

**Table 2.** Number and percent of sensitive and resistant nucleosomes within promoters at each time point following LPS stimulation. Nucleosomes with a log2 fold change in summit score greater than 1.5 combined with a p-value and FDR under 0.05 were classified as sensitive if more abundant in light digests and resistant if more abundant in heavy digests. All nucleosomes not meeting these thresholds were classified as neutral.

| time | # sensitive | # resistant | # neutral | % sensitive | % resistant |
| --- | --- | --- | --- | --- | --- |
| 0hr | 16857 | 22058 | 102667 | 11.9 | 15.6 |
| 5min | 21633 | 18521 | 104041 | 15.0 | 12.8 |
| 30min | 19344 | 15257 | 120513 | 12.5 | 9.8 |
| 1hr | 26045 | 21065 | 111904 | 16.4 | 13.2 |
| 4hr | 20235 | 17073 | 116513 | 13.2 | 11.1 |
| 12hr | 16841 | 12280 | 124942 | 10.9 | 8.0 |
| 24hr | 17287 | 14358 | 113488 | 11.9 | 9.9 |
| 48hr | 22848 | 12527 | 110927 | 15.6 | 8.6 |

Our final measure of nucleosome distribution, nucleosome occupancy, refers to alterations in the summit score of a nucleosome regardless of position on the DNA or width of the nucleosome peak. We defined alterations in nucleosome occupancy as having a log2 fold change greater than 5 with a p-value and FDR under 0.05 for any time point compared to the basal occupancy seen in 0hr (summary in Fig. 2E; counts in Table 1). By this definition, ∼5,000 nucleosomes at any time point showed increased (Fig. 2H, Supp. Table 3) or decreased (Fig. 2I, Supp. Table 4) occupancy following LPS stimulation. There is an inverted relationship between gains and losses in occupancy with early time points showing more promoters with increased occupancy and fewer promoters with decreased occupancy and later time points showing the opposite trend possibly indicating a recovery from earlier alterations(Fig. 2H-I). Unlike repositioning or fuzziness, the majority of nucleosomes with altered occupancies were common to all or more than three time points suggesting that while repositioning is transient, occupancy changes may be more durable. Promoters with increased occupancy at 5min and subsequent time points after LPS stimulation were enriched for RhoA and CDC42 activity which is important for macrophage chemotaxis and intravasation ^91,92^ (Supp. Fig. 3A). Other common pathways included IFN-γ, C-MYC, p38 via MAPK, and Ras signaling which were significantly enriched at most time points (Supp. Fig. 3A). Promoters with decreased occupancy were enriched for Hedgehog family, Ret tyrosine kinase, Reelin, NFAT, and nephrin signaling across most of the time points sampled (Supp. Fig. 2B). These pathways are associated with developmental processes or cancer and are only indirectly linked to macrophage activation or immune responses ^93–95^. This suggests that losses of nucleosome occupancy following LPS stimulation may be the result of nonspecific remodeling and are not necessarily relevant to cellular responses or expression of immune associated genes.

Our analysis of nucleosome distribution features indicates that nucleosome remodeling occurs over a discrete subset of promoters. Less than 10% of promoters had altered repositioning or fuzziness induced by LPS stimulation (Table 1). However, these promoters were frequently functionally relevant to macrophage response as indicated by enrichment of immune associated gene ontology terms. The repositioning or fuzziness alterations were predominantly transient in nature with minimal overlap between time points. In contrast, approximately 25% of promoters showed increased or decreased occupancy after LPS stimulation suggesting that this is a much more widespread effect of immune perturbation (Table 1). The majority of these alterations were found across multiple or all time points suggesting durable changes over time. Promoters with increased occupancy showed enrichments for macrophage associated ontology terms. However, the majority of genes with decreased occupancy were not functionally relevant to immune responses suggesting that these changes are non-specific and not directly related to cellular responses.Together this suggests that nucleosome repositioning, altered fuzziness, and increased occupancy, but not loss of occupancy, over specific promoters is important for proper cellular response to LPS stimulation.

### Alterations in nucleosome distribution occur over early and late LPS response genes and only partially associated with altered gene expression

Considering that nucleosome redistribution was limited to a subset of promoters associated with known immune functions, we expected that LPS response genes would have clear differences in promoter architecture associated with gene expression. We plotted nucleosome distribution over 234 LPS response genes (as defined by the cellular response to lipopolysaccharide gene ontology set GO:0071222) and a matched set of non-response genes (Fig. 3A). We found that response genes had a less pronounced nucleosome depleted region (NDR), similar upstream phasing, and more prominent -1 and +1 nucleosomes than matched genes in unstimulated macrophages (Fig. 3A). Five minutes after LPS stimulation, increased occupancy was observed over the -1 nucleosome and upstream phasing decreased in the response genes. Similar trends were observed until 24hr after stimulation when upstream phasing recovered in the response genes but the prominent -1 nucleosome maintained occupancy through the 48hr time point (Fig. 3A). These genes generally increased in expression over time with maximal expression at 1hr and 24hr after LPS stimulation (Fig. 3B). This result suggests that LPS response genes have a pre-established promoter architecture that is distinct from non-response genes and that dynamic alterations in nucleosome distribution occur early after and persist for days following stimulation. However, these promoter dynamics were not on the scale expected considering the extent of the immune response following LPS stimulation.This suggested that early and late LPS response genes are under separate regulatory control and should be evaluated independently.

**Figure 3.**
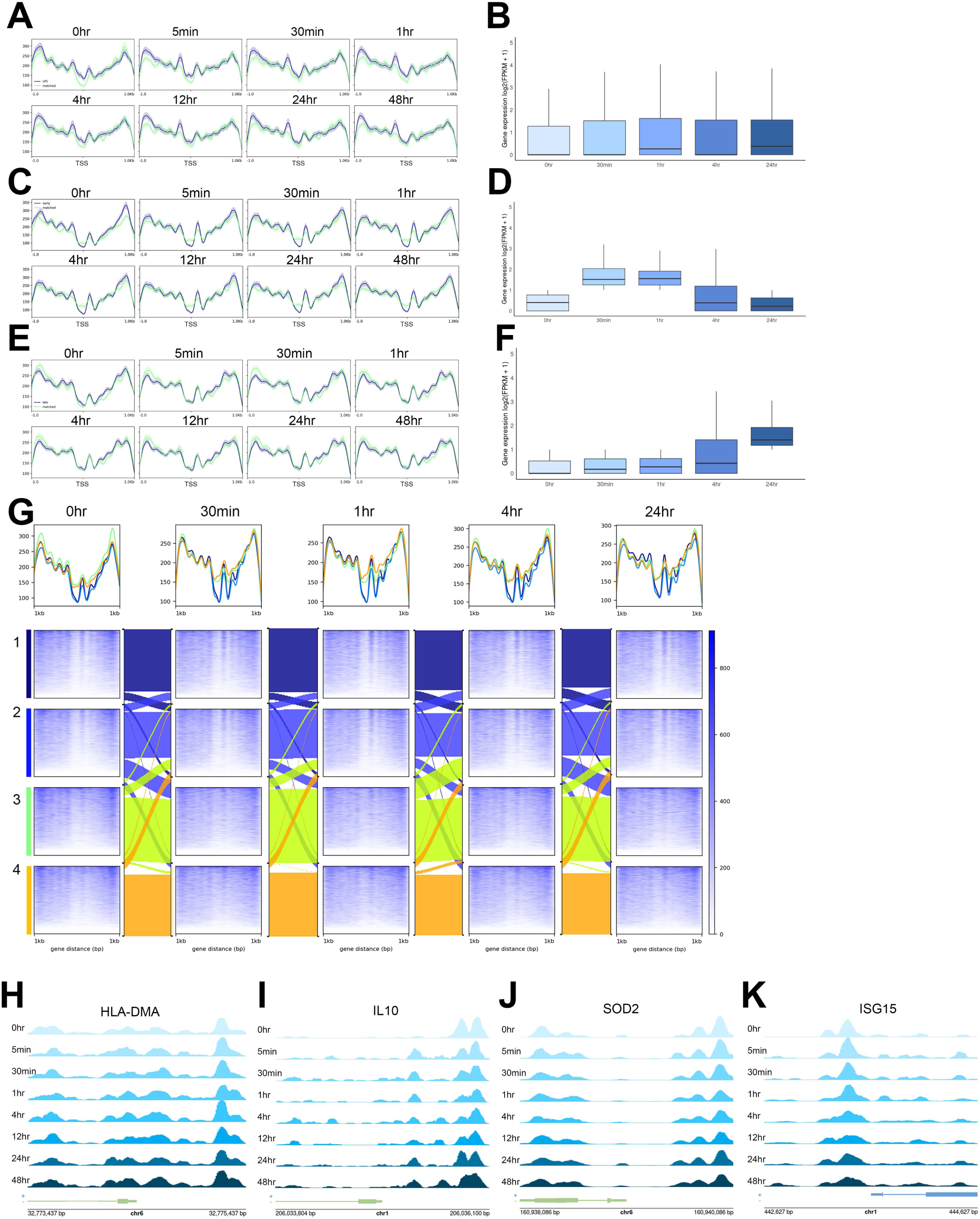
Nucleosome distribution is pre-established over early LPS response promoters in unstimulated cells. A. Line plots show average nucleosome distribution (solid line) with 95% confidence intervals (shadow) over GO defined LPS response and matched gene sets. B. Boxplots show gene expression (log2(FPKM+1)) for GO defined LPS response genes after stimulation. C. Line plots show average nucleosome distribution with 95% confidence intervals over early response genes as defined by expression profiles and matched gene sets. D. Boxplots show gene expression (log2(FPKM+1)) for expresion defined LPS early response genes after stimulation. E. Line plots show average nucleosome distribution with 95% confidence intervals over late response genes as defined by expression profiles and matched gene sets. F. Boxplots show gene expression (log2(FPKM+1)) for expresion defined LPS late response genes after stimulation. G. Heatmaps show nucleosome distribution over promoters grouped by gene expression quartiles at selected time points. Alluvial diagrams show changes in gene clustering between time points. H. Nucleosome distribution over the HLA-DMA promoter. I. Nucleosome distribution over the IL-10 promoter. J. Nucleosome distribution over the SOD2 promoter. K. Nucleosome distribution over the ISG15 promoter.

In order to determine whether early and late response genes have different basal promoter architectures and dynamic responses to LPS stimulation, we evaluated distribution patterns over two sets of LPS response promoters as defined by gene expression patterns with early genes expressed within one hour of stimulation and late genes expressed four hours after LPS stimulation. We first evaluated nucleosome distribution over a set of 257 curated LPS early response genes (Fig. 3C) which are rapidly expressed within 30min and 1hr following LPS stimulation but are not strongly expressed at later time points (Fig. 3D). We found that these primary LPS response genes had higher -1 nucleosome signal, more defined nucleosome phasing upstream of the promoter, and a more pronounced nucleosome depleted region over the immediate TSS compared to matched genes in unstimulated cells (Fig. 3C). These trends began to shift towards a stronger +1 nucleosome in the 5min and 30min samples and more defined downstream nucleosome positions 1 hour after LPS stimulation (Fig. 3C). Interestingly the altered phasing persisted throughout the time course with a consistent shift in periodicity between response and matched genes and a sharp +1 nucleosome present at all times in early genes. This result demonstrates that early response genes have a unique promoter architecture which is characterized by increased occupancy of a well-positioned +1 nucleosome immediately after LPS stimulation which persists up to 48 hours after stimulation.

We next evaluated the nucleosome distribution over a set of 516 late responding promoters (Fig. 3E) defined by minimal expression in unstimulated and early timepoints with greatest expression between 4-24hr after stimulation (Fig. 3F). We found that promoters in this gene set exhibited strikingly different nucleosome distribution patterns from the early LPS gene set. Late response genes lacked a highly occupied +1 nucleosome and had highly occupied but poorly positioned -2 and -1 nucleosomes in unstimulated cells (Fig. 3E). Increased occupancy over the +1 nucleosome was apparent within 5min of LPS stimulation but the overall distribution patterns were highly similar to the match gene set (Fig. 3E). The nucleosomes within these promoters were poorly positioned at all time points suggesting positional heterogeneity within the promoters.

Considering this heterogeneity, we wanted to evaluate the nucleosome profiles associated with highly expressed genes to determine if the profiles seen in selected gene sets corresponded with these patterns. We plotted nucleosome distribution over expression quartiles over time and found that highly expressed genes are defined by a pronounced nucleosome depleted region and a strongly positioned +1 nucleosome at all time points (Fig. 3G). This pattern is consistent with the profiles seen in the early LPS gene set (Fig. 3C). In contrast, weakly expressed or silent promoters lack a pronounced NDR and have a weakly positioned +1 nucleosome at all time points (Fig. 3G). This shows a clear association between nucleosome distribution patterns over promoters and gene expression which was consistent over time.

We expected that nucleosome remodeling events might precede gene expression alterations in a subset of genes while other remodeling events might not have a direct immediate effect on expression. We evaluated the gene expression profiles of promoters with altered nucleosome distribution defined by changes in position, fuzziness, and occupancy. Clustering expression patterns for redistributed nucleosomes at each time point showed that alterations in nucleosome distribution were not consistently associated with altered expression (Supp. Figs. 4-5). With changes in distribution occurring both in genes with static expression patterns and genes which were later under or overexpressed at later time points (Supp. Figs 4-5) indicating that nucleosome distribution alone is not sufficient for understanding gene expression changes. Additionally, we found that ∼30% of genes with altered nucleosome distribution within promoters were differentially expressed after LPS stimulation at any given time point (Supp. Table 5). This proportion was consistent over promoters defined by alterations in position, fuzziness, and occupancy and matched the genome wide average (Supp. Table 5) further indicating that altered distribution is not directly related to expression changes for the majority of genes.

The observed alterations in nucleosome distribution resulting from LPS stimulation are not necessarily related to alterations in gene expression. This lead us to propose that there are several distinct classes of promoters in terms of nucleosome distribution and gene expression that can be summarized as follows: (1) genes with no change in expression or distribution, (2) genes with no change in expression and altered nucleosome distribution, (3) genes with altered expression and no change in nucleosome distribution, and (4) genes with altered expression and nucleosome distribution. Each class of promoter is illustrated in the following examples. The HLA-DMA promoter, which encodes a MHC class II subunit, is consistently expressed at low levels and shows minimal nucleosome alterations after LPS stimulation (Fig. 3G). The antiinflammatory IL-10 promoter is not expressed in macrophages but gains nucleosome occupancy upstream of the TSS after LPS stimulation (Fig. 3H). In contrast, SOD2 is highly overexpressed in macrophages within 1 hour of LPS stimulation and shows minimal alterations in nucleosome distribution (Fig. 3I). The late responding ISG15 promoter is overexpressed 4 hours after LPS stimulation but shows increased signal over the +1 nucleosome within 5 minutes and a reduction in positioning over the -1 nucleosome 4 hours after LPS stimulation (Fig. 3J). These examples highlight the variation in nucleosome distribution profiles and show that promoter nucleosome distribution alone is not sufficient to explain alterations in gene expression following LPS stimulation. Similarly gene expression alone is not able to account for alterations in nucleosome distribution.

### Nucleosome sensitivity to MNase digestion over pol II promoters is highly dynamic following LPS stimulation

Considering that nucleosome distribution only partially explained alterations in gene expression following LPS stimulation at a subset of promoters, we next examined nucleosome sensitivity to MNase digestion. MNase sensitivity is determined by digesting fixed chromatin with a titration of MNase and comparing the resulting nucleosome maps to identify nucleosomes susceptible to overdigestion by MNase^58,59,61–63,83,96–98^. Under light-digestion conditions, a population of sensitive genomic regions is protected by nucleosomes, however, these footprints are lost under heavy digest conditions. Thus, a subset of nucleosomes displays a biochemical property rendering them susceptible to over digestion by MNase (schematic shown in Figure 1). We refer to nucleosomes which have greater coverage in light digests as “sensitive” to digestion and those which are preferentially released under heavier digest conditions as “resistant” to MNase digestion.

We identified MNase sensitive and resistant nucleosomes within 2kb of pol II promoters and used kmeans clustering to identify dominant sensitivity patterns as summarized in Figure 4. Clustering of promoters based on sensitivity patterns in unstimulated macrophages resulted in the identification of a class of diffusely sensitive group of promoters (Fig. 4A, 0hr cluster1) and several classes of promoters with highly positioned resistant promoters (Fig. 4A, 0hr clusters 2-7). These basal sensitivity patterns were dramatically disrupted within 5 minutes of LPS stimulation (Fig. 4A, 5min) and became increasingly more sensitive after 4 hours following LPS stimulation (Fig. 4A, 4hr-48hr, clusters 2-7). However, the diffusely sensitive cluster was largely maintained after LPS stimulation (Fig. 4A, all samples cluster 1) indicating that this class of promoter is more stable or less responsive to LPS induced alterations. This result suggests that, unlike nucleosome distribution, nucleosome sensitivity is highly dynamic and alterations in sensitivity occur at the majority of pol II promoters. This observation is quantitatively supported by Pearson’s correlation coefficients less than 0.4 between samples when comparing sensitivity signal in 10bp bins across promoters with the 12hr, 24hr, and 48hr samples clustering as outgroups (Fig. 3B). This grouping of time points corresponds to the expected timeframe for greatest chromatin remodeling as signaling cascades will rapidly activate early genes which is followed by a period of remodeling and expression of late responding genes several hours after initial stimulation ^18,20,28,40^.

**Figure 4.**
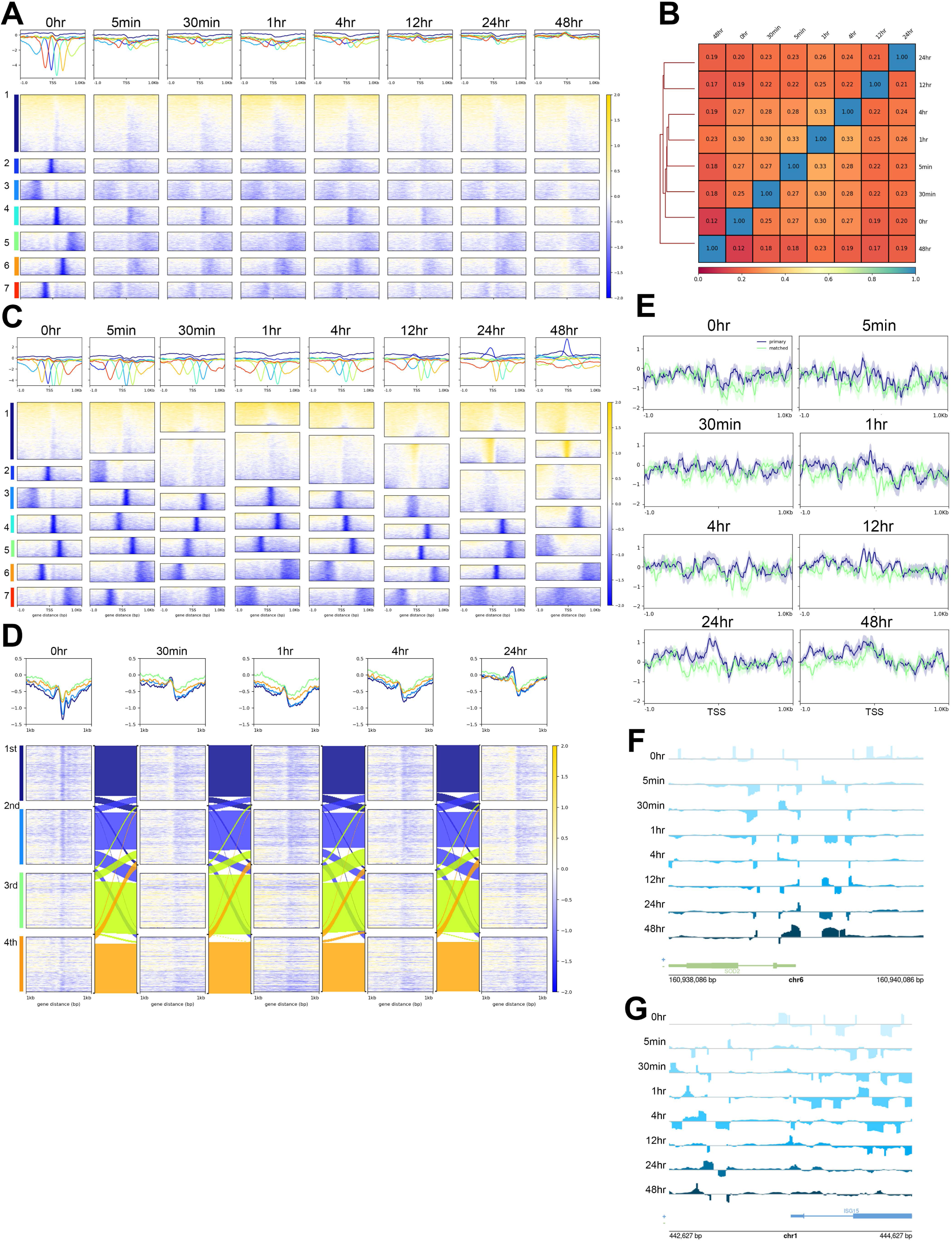
LPS stimulation results in highly dynamic alterations in nucleosome sensitivity across the majority of promoters. A. Heatmaps show nucleosome sensitivity centered on the TSSs of pol II genes. Sensitive regions are shown in yellow and resistant regions are shown in blue. Kmeans clustering was used to group genes based on unstimulated macrophages (0hr). Line plots above each plot show average signal values for each cluster. B. Pearson correlation scores for nucleosome sensitivity over promoters calculated in 10bp bins. C. Heatmaps show nucleosome sensitivity over all time points with kmeans clustering established independently for each time point. D. Nucleosome sensitivity plotted over the promoters of pol II genes separated into gene expression quartiles. E. Average nucleosome sensitivity profiles with 95% confidence intervals over primary LPS response and matched set of non-response genes. F. Nucleosome sensitivity profiles over the SOD2 promoter. G. Nucleosome sensitivity over the ISG15 promoter.

Since the basal macrophage nucleosome sensitivity pattern was not maintained after LPS stimulation, we were interested in further characterizing the sensitivity patterns that emerged at each time point. Accordingly, we independently generated kmeans clustered heatmaps of sensitivity signal over pol II promoters for each time point (Fig. 4C). This clustering produced a diffusely sensitive cluster and multiple positioned resistant clusters in all time points. This analysis showed a gradual and consistent increase in overall sensitivity between 5 minutes to 4 hours after LPS stimulation (Fig. 4C, 5 min-4 hour most apparent in cluster 2) and the emergence of a new type of cluster marked by a highly positioned sensitive nucleosome just upstream of the TSS starting 24 hours after stimulation (Fig. 4C, 12hr-48hr cluster 2). This trend of increasing sensitivity was supported by quantification of individual nucleosomes defined as either sensitive or resistant by a log2 fold change in summit score of at least 1.5 combined with a p-value and FDR under 0.05 (Table 2). Interestingly, the emergence of the unique position sensitivity cluster coincided with the timeframe for macrophage recovery after LPS stimulation following primary and secondary immune gene activation and may represent alterations at a set of promoters involved in recovery and polarization.

To investigate this, we performed gene ontology on promoters found in the highly positioned sensitive clusters in the 24hr and 48hr samples (Supp. Fig. 6). A total of 668 genes were common in this promoter class between the two time points and this included a number of macrophage associated genes including FCGR1A, FCGR2A, IL6R, and CCR7. These shared genes were enriched for biological process gene ontology terms related to translation, gene expression, and mRNA processing (Supp. Fig. 6A). The 2594 genes unique to the 24 hour cluster were enriched for similar ontology terms including mRNA processing, translation, tRNA modification, and ribosome biogenesis (Supp. Fig. 6B). In contrast, the 1576 genes unique to the 48 hour time point were enriched for ontology terms related to base excision repair, DNA replication, RNA processing, and DNA synthesis related to base repair (Supp. Fig. 6C). This final set of ontologies is of particular interest as TLR-4 stimulation results in production of reactive oxygen species which damage DNA thus necessitating repair as cells recover from stimulation.

We were interested in what nucleosome characteristics, if any, would distinguish the LPS induced positioned sensitive nucleosome at the 24hr and 48hr time points. We first plotted paired end fragment lengths for the light digests over a 2kb region centered on the TSS over promoters within individual sensitivity clusters for all time points (Supp. Fig. 7A-F). These plots show the distribution of fragment sizes as well as the density of fragments which shows nucleosome distribution, approximate linker length, and location of a nucleosome depleted region. We found that the strongly positioned resistant clusters had similar profiles over time (Supp. Fig. 7A-F, clusters 3-7). The diffusely sensitive cluster was similar over all samples with poorly defined nucleosomes and no discernable nucleosome depleted region over the immediate TSS (Supp. Fig. 7A-F, cluster 1). However, the highly positioned sensitive cluster showed an accumulation of subnucleosomal footprints between 100-140bp over the TSS in an otherwise nucleosome depleted region at 24hr and 48hr (Supp. Fig. 7E-F, cluster 2). This suggests that the highly localized sensitivity in these clusters may be due to non-nucleosome footprints, either in the form of a hexasome or a chromatin binding protein which protects the underlying DNA from MNase digestion.

We also evaluated the nucleosome positioning, reported as a fuzziness score, and occupancy, reported as a summit score, of nucleosomes underlying each independently generated sensitivity cluster using DANPOS3. We found that positioned sensitive clusters were more similar to positioned resistant clusters in terms of low fuzziness scores (Supp. Fig. 7G). However, the positioned sensitive promoters had high occupancy scores which were more similar to mixed or diffusely sensitive clusters (Supp. Fig. 7H). Together these characteristics define diffusely sensitive nucleosomes as being highly occupied but poorly positioned and lacking a nucleosome depleted region. Promoters with either positioned resistant or positioned sensitive regions are marked by a well defined NDR which is partially occupied by a subnucleosomal footprint in sensitive promoters.

Considering the functional relevance of genes identified by sensitivity clustering and the nucleosome profiles underlying each sensitivity cluster, we were interested in the relationship between nucleosome sensitivity and gene expression following LPS stimulation. Accordingly, we plotted nucleosome sensitivity over gene expression quartiles at 0hr, 30min, 1hr, 4hr, and 24hr following LPS treatment (Fig. 4D). Highly expressed genes at all time points were defined by a highly sensitive promoter in the -1 position and a highly resistant nucleosome in the +1 position (Fig. 4D, 1st quartile). In contrast, poorly expressed genes lacked the dynamic range between highly sensitive and highly resistant nucleosomes flanking the TSS with a weakly sensitive -1 and a weakly resistant +1 nucleosome (Fig. 4D, 4th quartile). In addition to these trends, there was also an overall increase in sensitivity in the -1 nucleosome at the 24 hour time point (Fig. 4D, 24hr 1st and 2nd quartiles). This same pattern of a sensitive -1 and resistant +1 nucleosome was also seen in primary LPS response gene promoters in unstimulated cells (Fig. 4E). However, this pattern was disrupted following LPS stimulation and was not appreciable until 24hr after stimulation. This applied to both LPS response and matched genes further suggesting that LPS derived sensitivity dynamics are widespread and impact the majority of genes regardless of LPS response status.

The dynamic nature of nucleosome sensitivity is apparent over individual promoters. The SOD2 promoter is diffusely sensitive in unstimulated macrophages, becomes largely resistant early after LPS stimulation, and develops a strongly sensitive region over the immediate TSS 48 hours after LPS stimulation (Fig. 4F). This gene is overexpressed within one hour of LPS stimulation and remains highly expressed 24 hours after stimulation. In contrast, the ISG15 promoter is initially highly resistant downstream of the TSS in unstimulated macrophages and trends towards a mixed sensitivity state at later time points when it is expressed in activated macrophages (Fig. 4G). These example loci show alterations in nucleosome sensitivity patterns with dynamic gains and losses of sensitivity across time points following LPS stimulation.

### Nucleosome sensitivity dynamics reflects alterations in active chromatin features following LPS stimulation

Next, we were interested in the functional relevance of sensitivity dynamics as related to active chromatin features and gene expression. Accordingly, we co-plotted the active H3K27ac mark along with ATACseq, which is a marker for accessible chromatin regions, with nucleosome distribution and sensitivity as summarized in Figure 5. We found that promoters with highly resistant positioned nucleosomes showed higher associations with the active H3K27ac mark and ATACseq in unstimulated macrophages (Fig. 5A, clusters 2-7). These genes had higher overall expression levels (Fig. 5F) and a more pronounced nucleosome depleted region around the TSS (Fig. 5A, clusters 2-7). In contrast, the diffusely sensitive cluster had weaker H3K27ac and ATACseq signal (Fig. 5A, cluster 1) and was poorly expressed (Fig. 5F). This association between positioned resistant nucleosomes and increased H3K27ac, ATACseq, and gene expression was maintained 30 minutes (Fig. 5B), 1 hour (Fig. 5C), and 4 hours (Fig. 5D) after LPS stimulation despite changes in sensitivity clustering patterns which resulted in the emergence of a promoter cluster which was largely neither sensitive nor resistant. This mixed cluster showed intermediate levels of active marks and gene expression (Fig. 5G-I). The highly positioned sensitive cluster which was unique to the 24 hour time point (Fig. 5E, cluster 2) was fundamentally different from the diffusely sensitive cluster seen in other time points. This cluster had the greatest H3K27ac and ATACseq signal and the highest level of gene expression (Fig. 5J) suggesting that the highly localized nature of this sensitivity pattern marks underlying promoters as fundamentally different from those which are marked by a diffuse sensitivity pattern.

**Figure 5.**
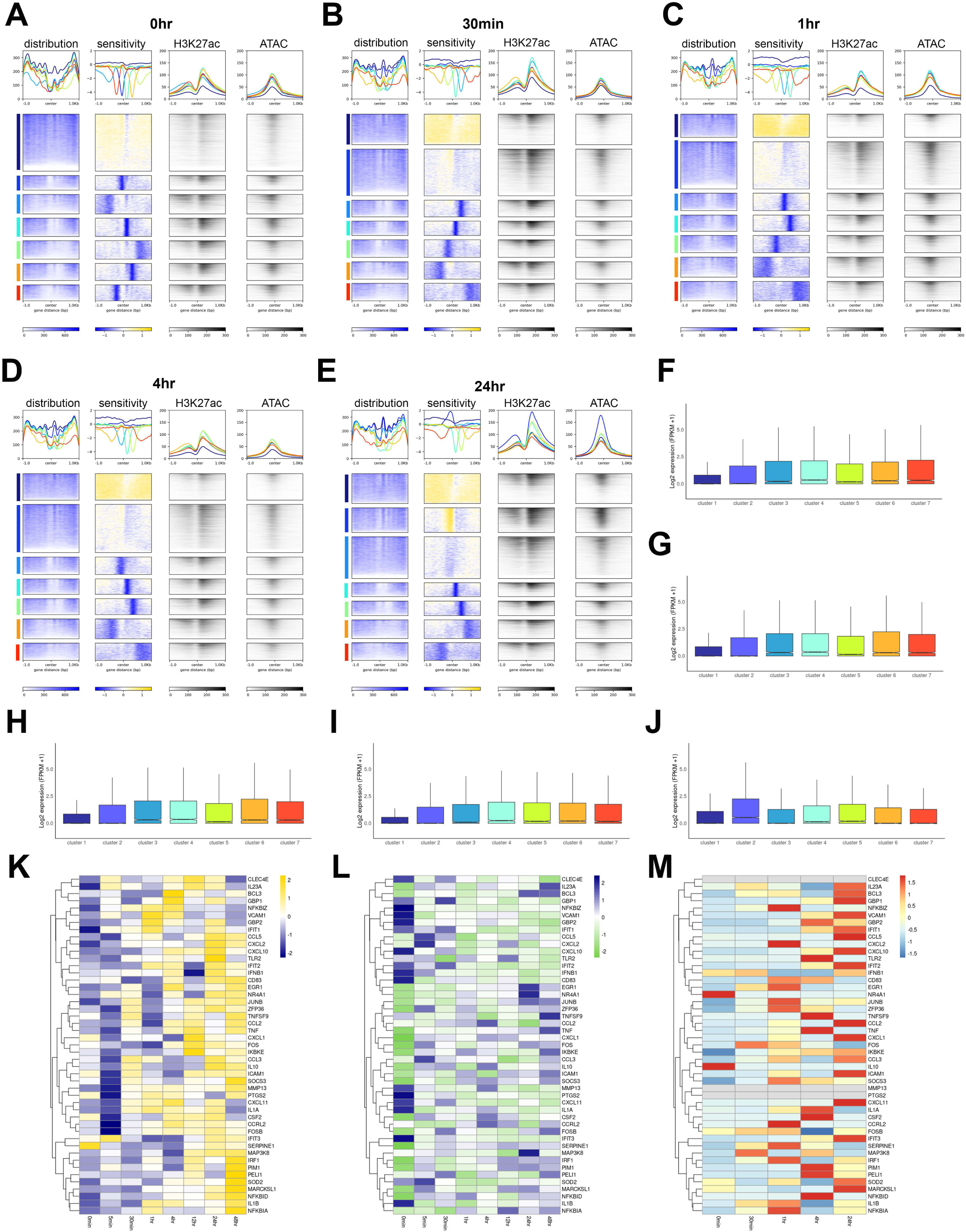
Nucleosome sensitivity is a robust marker of promoter activity throughout the macrophage response to LPS. A. Nucleosome distribution, sensitivity, H3K27ac ChIP, and ATAC signal over promoters in unstimulated macrophages clustered based on nucleosome sensitivity. B. Nucleosome distribution, sensitivity, H3K27ac ChIP, and ATAC signal over promoters in macrophages 30 minutes after LPS stimulation clustered based on nucleosome sensitivity. C. Nucleosome distribution, sensitivity, H3K27ac ChIP, and ATAC signal over promoters in macrophages 1 hour after LPS stimulation clustered based on nucleosome sensitivity. D. Nucleosome distribution, sensitivity, H3K27ac ChIP, and ATAC signal over promoters in macrophages 4 hours after LPS stimulation clustered based on nucleosome sensitivity. E. Nucleosome distribution, sensitivity, H3K27ac ChIP, and ATAC signal over promoters in macrophages 24 hours after LPS stimulation clustered based on nucleosome sensitivity. F. Gene expression (log2(FPKM+1)) for each sensitivity cluster in unstimulated macrophages. The median expression level per cluster is shown as a black line across the boxplot. G. Gene expression (log2(FPKM+1)) for each sensitivity cluster in macrophages 30 minutes after LPS stimulation. The median expression level per cluster is shown as a black line across the boxplot. H. Gene expression (log2(FPKM+1)) for each sensitivity cluster in macrophages one hour after LPS stimulation. The median expression level per cluster is shown as a black line across the boxplot. I. Gene expression (log2(FPKM+1)) for each sensitivity cluster in macrophages 4 hours after LPS stimulation. The median expression level per cluster is shown as a black line across the boxplot. Gene expression (log2(FPKM+1)) for each sensitivity cluster in macrophages 24 hours after LPS stimulation. The median expression level per cluster is shown as a black line across the boxplot. K. Nucleosome sensitivity z scores across promoters of primary LPS response genes. Sensitive promoters are shown in gold and resistant promoters are shown in blue. L. Nucleosome occupancy z scores over the same response genes with highest occupancy in blue and lowest in green. M. Gene expression z scores for the same set of genes with highest expression in red.

Considering the relationship between positioned resistant nucleosomes and active promoters, we next directly compared sensitivity profiles (Fig. 5K), nucleosome distribution (Fig. 5L), and gene expression (Fig. 5M) over individual LPS primary response promoters. We found that the majority of promoters in this set are initially resistant for the first hour following LPS stimulation and trend towards increasing sensitivity between 12-48hr following exposure (Fig. 5K). Nucleosome sensitivity was not directly related to nucleosome occupancy with both sensitive and resistant promoters showing mixed occupancy states (Fig. 5L). High gene expression (Fig. 5M) was associated with a sensitive promoter state preceding the time of expression (e.g. IL23A, second from top, showing highest sensitivity at 12hr and highest expression at 24hr). Together this indicates that nucleosome sensitivity is intrinsically related to promoter activity with diffuse sensitivity marking silent genes and positioned sensitive or resistant nucleosomes marking active genes.

These general observations led us to closely examine promoter features of individual LPS response genes. We plotted nucleosome distribution, sensitivity, ATAC, and H3K27ac signal over the IL1B, ICAM-1, NFKBIA, and IFIT3 promoters (Fig. 6A). IL1B and NFKBIA are early response genes with maximum expression 1hr after LPS stimulation and later activation at 24hr. ICAM-1 and IFIT3 are late genes with limited expresion before 24hr (Fig. 6B). Each of these genes shows distinctive promoter dynamics with early alterations in nucleosome positioning and occupancy, highly dynamic sensitivity profiles, and an accumulation of ATAC and H3K27ac corresponding with expression. The high temporal resolution shows a loss of nucleosome occupancy over IL1B within 5 minutes of LPS stimulation and a redistribution of sensitivity to the gene body within the first 30 minutes which corresponded to an increase in ATAC signal and maximum gene expression at 1hr. While H3K27ac is associated with this promoter, the signal accumulated gradually and was most prominent 24hr after LPS stimulation rather than at the point of highest expression. ICAM-1 loses a highly occupied -1 nucleosome, becomes less resistant to MNsae digestion, and gains ATAC and H3K27ac signal prior to maximum expression at 24hr after LPS stimulation. NFKBIA maintains a strong nucleosome depleted region and has increased +1 nucleosome occupancy starting 5 minutes after LPS stimulation. This promoter is highly sensitive over the inmate TSS and strong ATAC and H3K27ac signal through the time course. IFIT3 loses occupancy across the promoter and shows transient resistance patterns prior to maximum expression at 24hr. These promoters represent the dynamic nature of promoter architecture following LPS stimulation over early and late response genes.

**Figure 6.**
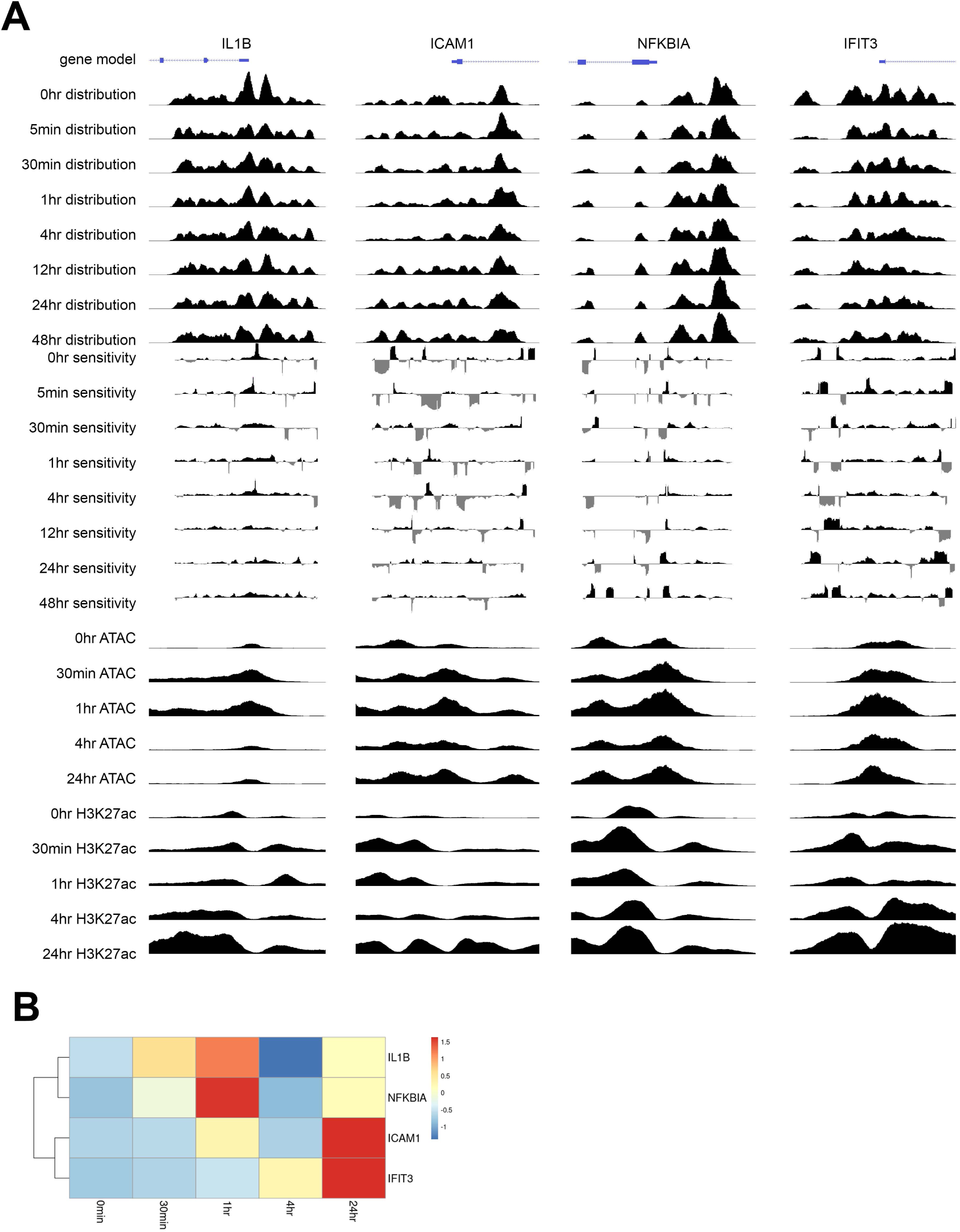
Altered promoter architecture over immune genes induced by LPS stimulation occurs prior to alterations in gene expression. A. Nucleosome distribution, nucleosome sensitivity, ATAC signal, and H3K27ac signal over selected LPS response gene promoters. Each promoter is shown in a 2kb region and signal values for all datatypes have set axes over time. B. Heatmap shows z scores of gene expression over time for the promoters in A. Peak IL1B expression occurs at 30min-1hr while NFKBIA is highly expressed at 1hr. ICAM1 is moderately expressed at 1hr and is highly expressed at 24hr and IFIT3 is highly expressed 24hr after stimulation with LPS.

## Discussion

In this study, we comprehensively identified promoters with altered nucleosome distribution and sensitivity patterns in macrophages following LPS stimulation. We found that approximately 10% of pol II promoters are rapidly remodeled in a time dependent and often transient manner (Fig. 2). Early remodeled promoters were enriched for immune-related genes, including key promoters within the ISG and JAK/STAT pathways, as revealed by gene ontology analysis. In contrast, remodeled promoters at later time points were enriched for more general terms (Supp. Fig. 1). Additionally, a subset of LPS response promoters were poised for rapid expression in unstimulated cells and did not undergo remodeling after LPS stimulation (Fig. 3). While nucleosome distribution alterations were highly specific and limited to a subset of promoters, alterations in nucleosome sensitivity were widespread and did not return to a basal state (Fig. 4). Active promoters with highly positioned resistant nucleosomes became increasingly more sensitive after LPS stimulation and a new class of hyper-active positioned sensitive nucleosomes emerged 24hr and 48hr after LPS stimulation (Fig. 5). Together these results show that both nucleosome distribution and sensitivity states are important for appropriate temporal regulation of gene expression following immune stimulation. This provides a framework for temporal dynamics underpinning innate immune responses and any other temporally regulated processes within the nucleus.

Our observations of limited and specific remodeling of selected promoters is consistent with previous work with reactivation of Kaposi’s Sarcoma Associated Herpesvirus and LPS stimulation where a subset of promoters were transiently remodeled following stimulation ^40,54,55^. However this differs from observations that lymphoblastoid cells undergo limited remodeling immediately following stimulation with heat-killed *Salmonella^60^*. This discrepancy may result from phenotypic differences between cell types, methodological differences, and time point selection as TLR-4 induction is similar in LPS and heat killed bacteria^99^. The temporal resolution of remodeling events observed in this study supports inferences of remodeling made through analysis of ATAC accessible regions following LPS stimulation ^33,35^. Additionally, the specific and targeted remodeling response further supports the observation that nucleosome distribution is highly similar in disparate cell types and during cancer development ^56,61^ as a potent stimulus along with highly refined time-resolved observations are necessary to capture transient remodeling events.

Our analysis of nucleosome sensitivity provides new insights into macrophage responses to LPS stimulation. We found highly dynamic responses across the majority of pol II promoters following stimulation, which echoed widespread alterations seen in other systems ^58–61^. The general trend for increasing sensitivity over promoters was also seen in lymphoblastoid cells treated with heat-killed *Salmonella* suggesting that this might be a common feature following immune insult ^60,61^. The association between positioned resistant nucleosomes and promoter activity was maintained over time even with the emergence of a unique highly sensitive positioned cluster of promoters which was only observed at the latest time points. The hyperactivity of these genes was associated with protection from a subnucleosomal structure suggesting hexasome occupancy or regulatory factor binding. This pattern has previously been observed in H2A.Z depletion in breast epithelial cells suggesting possible involvement of histone variants in appropriate gene regulatory programs ^61^.

In conclusion, our study elucidates the intricate dynamics of nucleosome distribution and sensitivity in macrophages following LPS stimulation. We observed that nucleosome remodeling and sensitivity changes are both time-dependent and gene-specific, highlighting the complexity of chromatin-based gene regulation during immune responses. Our findings suggest a model where early LPS response genes undergo rapid and transient nucleosome remodeling, while late response genes exhibit sustained sensitivity changes, potentially involving subnucleosomal structures or regulatory factor binding. This dual mechanism of nucleosome dynamics ensures precise temporal regulation of gene expression, enabling macrophages to effectively respond to external stimuli (Fig. 7). By providing a comprehensive framework for understanding these temporal chromatin changes, our study advances the field of macrophage cell biology and offers insights applicable to other cellular contexts. Future research should focus on dissecting the molecular mechanisms driving these nucleosome alterations and their broader implications for gene regulation and cellular function.

## ACKNOWLEDGEMENTS

We thank Lorea Arambarri and Shahin Behrouz Sharif for constructive conversations regarding data analysis and all members of the Dennis and Bass labs for critiquing the manuscript.

## AUTHOR CONTRIBUTIONS

JMB: Conceptualization, Methodology, Formal Analysis, Visualization, Software, Data curation, Writing--original draft and editing. BDB: Conceptualization, Methodology, Writing—review. MK: Writing—review. HWB: Methodology, Writing—review and editing. JHD: Funding acquisition, Conceptualization, Supervision, Resources, Writing – review & editing.

## DATA AVAILABILITY

All raw and processed sequencing data generated in this study have been submitted to the NCBI Gene Expression Omnibus (GEO; https://www.ncbi.nlm.nih.gov/geo/) under accession number GSE279622.

---

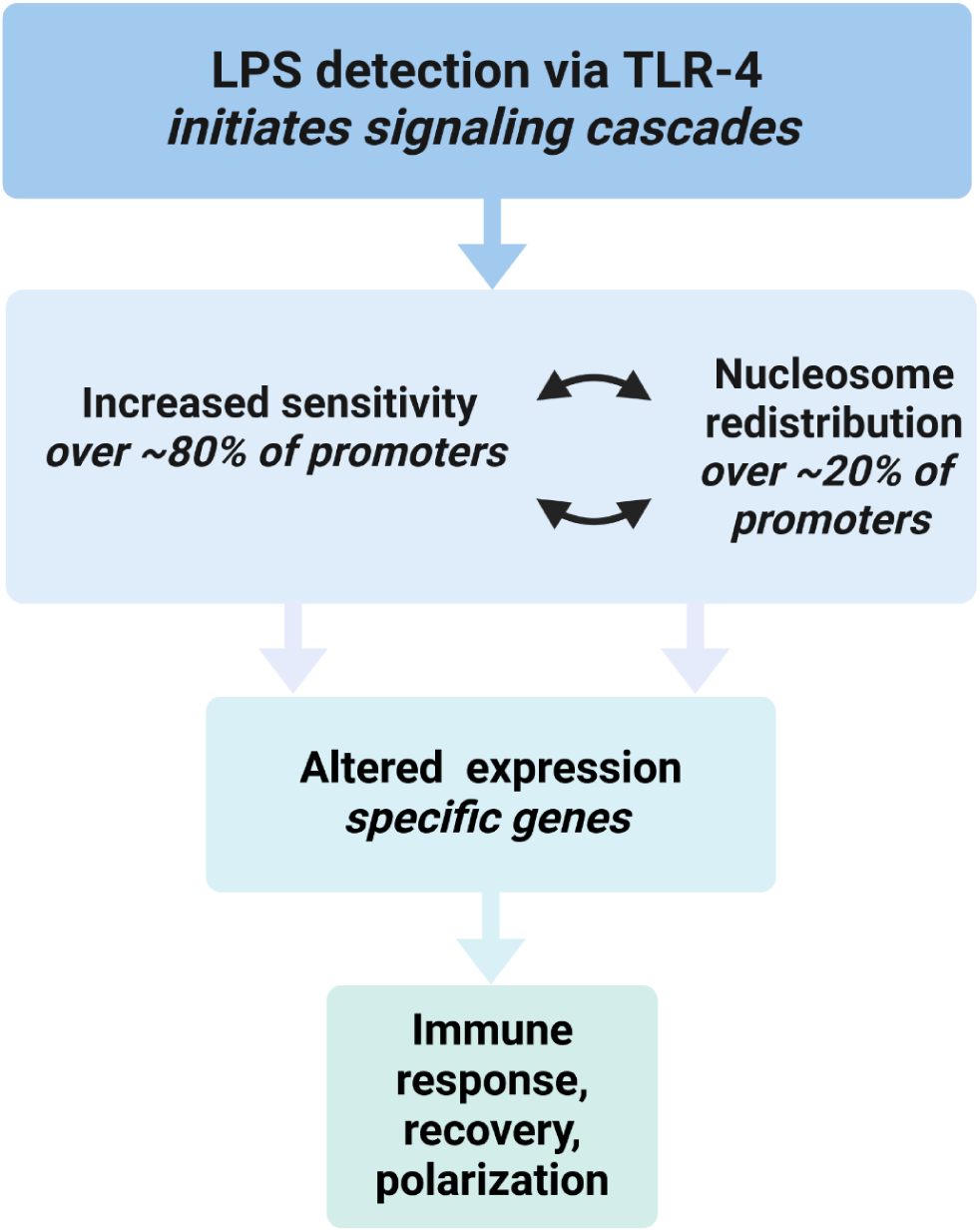
Summary figure. Macrophage detection of LPS occurs via TLR-4 and results in signaling cascades leading to rapid translocation of transcription factors to the nucleus. The majority of promoters become increasingly sensitive to MNase digestion in the minutes and hours following LPS stimulation while nucleosomes within a subset of promoters are redistributed. Increased expression of a select number of genes, some of which have undergone chromatin remodeling, results in appropriate immune responses, recovery, and polarization.

