## Supplementary material for "Dynamic nucleosome redistribution and increases in nucleosome sensitivity underpin THP-1 macrophage response to LPS": supp_figures

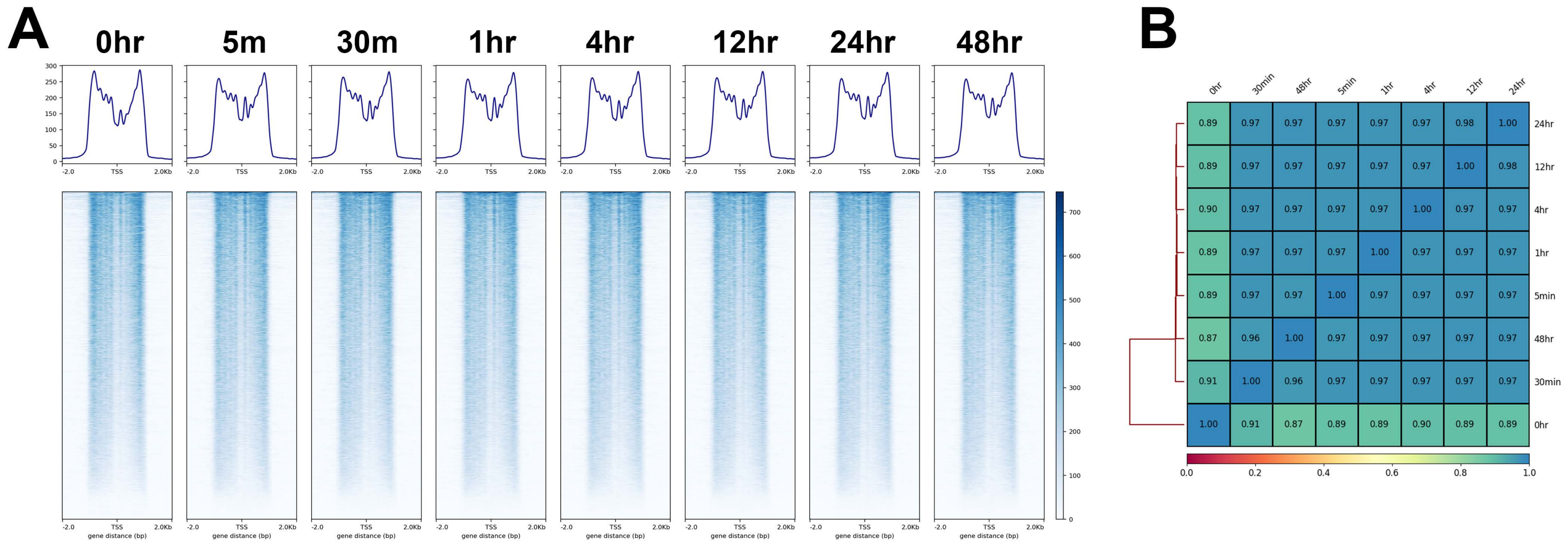
A
0hr
5m
30m
1hr
4hr
12hr
24hr
48hr
300
250
200
150
100
50
0
-2.0
TSS
2.0Kb
gene distance (bp)
700
600
500
400
300
200
100
0
B
0hr
30min
48hr
5min
1hr
4hr
12hr
24hr
0.89 0.97 0.97 0.97 0.97 0.97 0.98 1.00 24hr
0.89 0.97 0.97 0.97 0.97 0.97 1.00 0.98 12hr
0.90 0.97 0.97 0.97 0.97 1.00 0.97 0.97 4hr
0.89 0.97 0.97 0.97 1.00 0.97 0.97 0.97 1hr
0.89 0.97 0.97 1.00 0.97 0.97 0.97 0.97 5min
0.87 0.96 1.00 0.97 0.97 0.97 0.97 0.97 48hr
0.91 1.00 0.96 0.97 0.97 0.97 0.97 0.97 30min
1.00 0.91 0.87 0.89 0.89 0.90 0.89 0.89 0hr
0.0
0.2
0.4
0.6
0.8
1.0

#### Supp. Figure 1. Nucleosome distribution over pol II promoters

A. Heatmaps show nucleosome distribution over a 4kb window centered on pol II TSSs. Line plots above each heatmap show the average nucleosome distribution profile. B. Pearson's correlation of nucleosome signal in 10bp bin over promoters across time points. Correlation values are reported in each square.

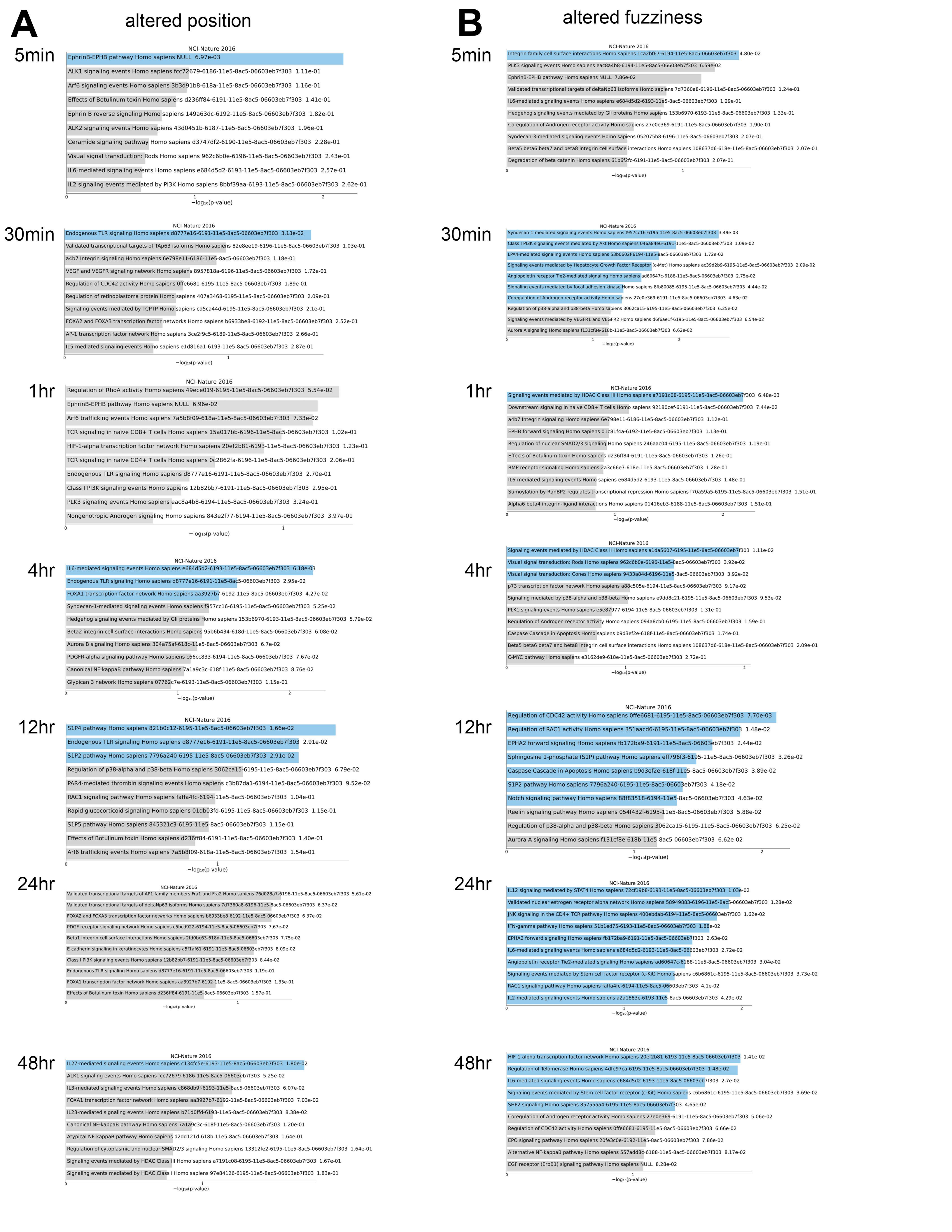
A
altered position
5min
NCI-Nature 2016
EphrinB-EPHB pathway Homo sapiens NULL 6.97e-03
ALK1 signaling events Homo sapiens fcc72679-6186-11e5-8ac5-06603eb7f303 1.11e-01
Arf6 signaling events Homo sapiens 3b3d91b8-618a-11e5-8ac5-06603eb7f303 1.16e-01
Effects of Botulinum toxin Homo sapiens d236ff84-6191-11e5-8ac5-06603eb7f303 1.41e-01
Ephrin B reverse signaling Homo sapiens 149a63dc-6192-11e5-8ac5-06603eb7f303 1.82e-01
ALK2 signaling events Homo sapiens 43d0451b-6187-11e5-8ac5-06603eb7f303 1.96e-01
Ceramide signaling pathway Homo sapiens d3747df2-6190-11e5-8ac5-06603eb7f303 2.28e-01
Visual signal transduction: Rods Homo sapiens 962c6b0e-6196-11e5-8ac5-06603eb7f303 2.43e-01
IL6-mediated signaling events Homo sapiens e684d5d2-6193-11e5-8ac5-06603eb7f303 2.57e-01
IL2 signaling events mediated by PI3K Homo sapiens 8bbf39aa-6193-11e5-8ac5-06603eb7f303 2.62e-01
−log10(p-value)
30min
NCI-Nature 2016
Endogenous TLR signaling Homo sapiens d8777e16-6191-11e5-8ac5-06603eb7f303 3.13e-02
Validated transcriptional targets of TAp63 isoforms Homo sapiens 82e8ee19-6196-11e5-8ac5-06603eb7f303 1.03e-01
a4b7 Integrin signaling Homo sapiens 6e798e11-6186-11e5-8ac5-06603eb7f303 1.18e-01
VEGF and VEGFR signaling network Homo sapiens 8957818a-6196-11e5-8ac5-06603eb7f303 1.72e-01
Regulation of CDC42 activity Homo sapiens 0ffe6681-6195-11e5-8ac5-06603eb7f303 1.89e-01
Regulation of retinoblastoma protein Homo sapiens 407a3468-6195-11e5-8ac5-06603eb7f303 2.09e-01
Signaling events mediated by TCPTP Homo sapiens cd5ca44d-6195-11e5-8ac5-06603eb7f303 2.1e-01
FOXA2 and FOXA3 transcription factor networks Homo sapiens b6933be8-6192-11e5-8ac5-06603eb7f303 2.52e-01
AP-1 transcription factor network Homo sapiens 3ce2f9c5-6189-11e5-8ac5-06603eb7f303 2.66e-01
IL5-mediated signaling events Homo sapiens e1d816a1-6193-11e5-8ac5-06603eb7f303 2.87e-01
−log10(p-value)
1hr
NCI-Nature 2016
Regulation of RhoA activity Homo sapiens 49ece019-6195-11e5-8ac5-06603eb7f303 5.54e-02
EphrinB-EPHB pathway Homo sapiens NULL 6.96e-02
Arf6 trafficking events Homo sapiens 7a5b8f09-618a-11e5-8ac5-06603eb7f303 7.33e-02
TCR signaling in naive CD8+ T cells Homo sapiens 15a017bb-6196-11e5-8ac5-06603eb7f303 1.02e-01
HIF-1-alpha transcription factor network Homo sapiens 20ef2b81-6193-11e5-8ac5-06603eb7f303 1.23e-01
TCR signaling in naive CD4+ T cells Homo sapiens 0c2862fa-6196-11e5-8ac5-06603eb7f303 2.06e-01
Endogenous TLR signaling Homo sapiens d8777e16-6191-11e5-8ac5-06603eb7f303 2.70e-01
Class I PI3K signaling events Homo sapiens 12b82bb7-6191-11e5-8ac5-06603eb7f303 2.95e-01
PLK3 signaling events Homo sapiens eac8a4b8-6194-11e5-8ac5-06603eb7f303 3.24e-01
Nongenotropic Androgen signaling Homo sapiens 843e2f77-6194-11e5-8ac5-06603eb7f303 3.97e-01
−log10(p-value)
4hr
NCI-Nature 2016
IL6-mediated signaling events Homo sapiens e684d5d2-6193-11e5-8ac5-06603eb7f303 6.18e-03
Endogenous TLR signaling Homo sapiens d8777e16-6191-11e5-8ac5-06603eb7f303 2.95e-02
FOXA1 transcription factor network Homo sapiens aa3927b7-6192-11e5-8ac5-06603eb7f303 4.27e-02
Syndecan-1-mediated signaling events Homo sapiens f957cc16-6195-11e5-8ac5-06603eb7f303 5.25e-02
Hedgehog signaling events mediated by Gli proteins Homo sapiens 153b6970-6193-11e5-8ac5-06603eb7f303 5.79e-02
Beta2 integrin cell surface interactions Homo sapiens 95b6b434-618d-11e5-8ac5-06603eb7f303 6.08e-02
Aurora B signaling Homo sapiens 304a75af-618c-11e5-8ac5-06603eb7f303 6.7e-02
PDGFR-alpha signaling pathway Homo sapiens c66cc833-6194-11e5-8ac5-06603eb7f303 7.67e-02
Canonical NF-kappaB pathway Homo sapiens 7a1a9c3c-618f-11e5-8ac5-06603eb7f303 8.76e-02
Glypican 3 network Homo sapiens 07762c7e-6193-11e5-8ac5-06603eb7f303 1.15e-01
−log10(p-value)
12hr
NCI-Nature 2016
S1P4 pathway Homo sapiens 821b0c12-6195-11e5-8ac5-06603eb7f303 1.66e-02
Endogenous TLR signaling Homo sapiens d8777e16-6191-11e5-8ac5-06603eb7f303 2.91e-02
S1P2 pathway Homo sapiens 7796a240-6195-11e5-8ac5-06603eb7f303 2.91e-02
Regulation of p38-alpha and p38-beta Homo sapiens 3062ca15-6195-11e5-8ac5-06603eb7f303 6.79e-02
PAR4-mediated thrombin signaling events Homo sapiens c3b87da1-6194-11e5-8ac5-06603eb7f303 9.52e-02
RAC1 signaling pathway Homo sapiens faffa4fc-6194-11e5-8ac5-06603eb7f303 1.04e-01
Rapid glucocorticoid signaling Homo sapiens 01db03fd-6195-11e5-8ac5-06603eb7f303 1.15e-01
S1P5 pathway Homo sapiens 845321c3-6195-11e5-8ac5-06603eb7f303 1.15e-01
Effects of Botulinum toxin Homo sapiens d236ff84-6191-11e5-8ac5-06603eb7f303 1.40e-01
Arf6 trafficking events Homo sapiens 7a5b8f09-618a-11e5-8ac5-06603eb7f303 1.54e-01
−log10(p-value)
24hr
NCI-Nature 2016
Validated transcriptional targets of AP1 family members Fra1 and Fra2 Homo sapiens 76d028a7-6196-11e5-8ac5-06603eb7f303 5.61e-02
Validated transcriptional targets of deltaNp63 isoforms Homo sapiens 7d7360a8-6196-11e5-8ac5-06603eb7f303 6.37e-02
FOXA2 and FOXA3 transcription factor networks Homo sapiens b6933be8-6192-11e5-8ac5-06603eb7f303 6.37e-02
PDGF receptor signaling network Homo sapiens c5bcd922-6194-11e5-8ac5-06603eb7f303 7.67e-02
Beta1 integrin cell surface interactions Homo sapiens 2fd0bc63-618d-11e5-8ac5-06603eb7f303 7.75e-02
E-cadherin signaling in keratinocytes Homo sapiens a5f1af61-6191-11e5-8ac5-06603eb7f303 8.09e-02
Class I PI3K signaling events Homo sapiens 12b82bb7-6191-11e5-8ac5-06603eb7f303 8.44e-02
Endogenous TLR signaling Homo sapiens d8777e16-6191-11e5-8ac5-06603eb7f303 1.19e-01
FOXA1 transcription factor network Homo sapiens aa3927b7-6192-11e5-8ac5-06603eb7f303 1.35e-01
Effects of Botulinum toxin Homo sapiens d236ff84-6191-11e5-8ac5-06603eb7f303 1.57e-01
−log10(p-value)
48hr
NCI-Nature 2016
IL27-mediated signaling events Homo sapiens c134fc5e-6193-11e5-8ac5-06603eb7f303 1.80e-02
ALK1 signaling events Homo sapiens fcc72679-6186-11e5-8ac5-06603eb7f303 5.25e-02
IL3-mediated signaling events Homo sapiens c868db9f-6193-11e5-8ac5-06603eb7f303 6.07e-02
FOXA1 transcription factor network Homo sapiens aa3927b7-6192-11e5-8ac5-06603eb7f303 7.03e-02
IL23-mediated signaling events Homo sapiens b71d0ffd-6193-11e5-8ac5-06603eb7f303 8.38e-02
Canonical NF-kappaB pathway Homo sapiens 7a1a9c3c-618f-11e5-8ac5-06603eb7f303 1.20e-01
Atypical NF-kappaB pathway Homo sapiens d2dd121d-618b-11e5-8ac5-06603eb7f303 1.64e-01
Regulation of cytoplasmic and nuclear SMAD2/3 signaling Homo sapiens 13312fe2-6195-11e5-8ac5-06603eb7f303 1.64e-01
Signaling events mediated by HDAC Class III Homo sapiens a7191c08-6195-11e5-8ac5-06603eb7f303 1.67e-01
Signaling events mediated by HDAC Class I Homo sapiens 97e84126-6195-11e5-8ac5-06603eb7f303 1.83e-01
−log10(p-value)
B
altered fuzziness
5min
NCI-Nature 2016
Integrin family cell surface interactions Homo sapiens 1ca2bf67-6194-11e5-8ac5-06603eb7f303 4.80e-02
PLK3 signaling events Homo sapiens eac8a4b8-6194-11e5-8ac5-06603eb7f303 6.59e-02
EphrinB-EPHB pathway Homo sapiens NULL 7.86e-02
Validated transcriptional targets of deltaNp63 isoforms Homo sapiens 7d7360a8-6196-11e5-8ac5-06603eb7f303 1.24e-01
IL6-mediated signaling events Homo sapiens e684d5d2-6193-11e5-8ac5-06603eb7f303 1.29e-01
Hedgehog signaling events mediated by Gli proteins Homo sapiens 153b6970-6193-11e5-8ac5-06603eb7f303 1.33e-01
Coregulation of Androgen receptor activity Homo sapiens 27e0e369-6191-11e5-8ac5-06603eb7f303 1.90e-01
Syndecan-3-mediated signaling events Homo sapiens 052075b8-6196-11e5-8ac5-06603eb7f303 2.07e-01
Beta5 beta6 beta7 and beta8 integrin cell surface interactions Homo sapiens 108637d6-618e-11e5-8ac5-06603eb7f303 2.07e-01
Degradation of beta catenin Homo sapiens 61b6f2fc-6191-11e5-8ac5-06603eb7f303 2.07e-01
−log10(p-value)
30min
NCI-Nature 2016
Syndecan-1-mediated signaling events Homo sapiens f957cc16-6195-11e5-8ac5-06603eb7f303 3.49e-03
Class I PI3K signaling events mediated by Akt Homo sapiens 046a84e6-6191-11e5-8ac5-06603eb7f303 1.09e-02
LPA4-mediated signaling events Homo sapiens 53b0602f-6194-11e5-8ac5-06603eb7f303 1.72e-02
Signaling events mediated by Hepatocyte Growth Factor Receptor (c-Met) Homo sapiens ac39d2b9-6195-11e5-8ac5-06603eb7f303 2.09e-02
Angiopoietin receptor Tie2-mediated signaling Homo sapiens ad60647c-6188-11e5-8ac5-06603eb7f303 2.75e-02
Signaling events mediated by focal adhesion kinase Homo sapiens 8fb80085-6195-11e5-8ac5-06603eb7f303 4.44e-02
Coregulation of Androgen receptor activity Homo sapiens 27e0e369-6191-11e5-8ac5-06603eb7f303 4.63e-02
Regulation of p38-alpha and p38-beta Homo sapiens 3062ca15-6195-11e5-8ac5-06603eb7f303 6.25e-02
Signaling events mediated by VEGFR1 and VEGFR2 Homo sapiens d6f6ae1f-6195-11e5-8ac5-06603eb7f303 6.54e-02
Aurora A signaling Homo sapiens f131cf8e-618b-11e5-8ac5-06603eb7f303 6.62e-02
−log10(p-value)
1hr
NCI-Nature 2016
Signaling events mediated by HDAC Class III Homo sapiens a7191c08-6195-11e5-8ac5-06603eb7f303 6.48e-03
Downstream signaling in naive CD8+ T cells Homo sapiens 92180cef-6191-11e5-8ac5-06603eb7f303 7.44e-02
a4b7 Integrin signaling Homo sapiens 6e798e11-6186-11e5-8ac5-06603eb7f303 1.12e-01
EPHB forward signaling Homo sapiens 01c81f4a-6192-11e5-8ac5-06603eb7f303 1.13e-01
Regulation of nuclear SMAD2/3 signaling Homo sapiens 246aac04-6195-11e5-8ac5-06603eb7f303 1.19e-01
Effects of Botulinum toxin Homo sapiens d236ff84-6191-11e5-8ac5-06603eb7f303 1.26e-01
BMP receptor signaling Homo sapiens 2a3c66e7-618e-11e5-8ac5-06603eb7f303 1.28e-01
IL6-mediated signaling events Homo sapiens e684d5d2-6193-11e5-8ac5-06603eb7f303 1.48e-01
Sumoylation by RanBP2 regulates transcriptional repression Homo sapiens f70a59a5-6195-11e5-8ac5-06603eb7f303 1.51e-01
Alpha6 beta4 integrin-ligand interactions Homo sapiens 01416eb3-6188-11e5-8ac5-06603eb7f303 1.51e-01
−log10(p-value)
4hr
NCI-Nature 2016
Signaling events mediated by HDAC Class II Homo sapiens a1da5607-6195-11e5-8ac5-06603eb7f303 1.11e-02
Visual signal transduction: Rods Homo sapiens 962c6b0e-6196-11e5-8ac5-06603eb7f303 3.92e-02
Visual signal transduction: Cones Homo sapiens 9433a84d-6196-11e5-8ac5-06603eb7f303 3.92e-02
p73 transcription factor network Homo sapiens a88c505e-6194-11e5-8ac5-06603eb7f303 9.17e-02
Signaling mediated by p38-alpha and p38-beta Homo sapiens e9dd8c21-6195-11e5-8ac5-06603eb7f303 9.53e-02
PLK1 signaling events Homo sapiens e5e87977-6194-11e5-8ac5-06603eb7f303 1.31e-01
Regulation of Androgen receptor activity Homo sapiens 094a8cb0-6195-11e5-8ac5-06603eb7f303 1.59e-01
Caspase Cascade in Apoptosis Homo sapiens b9d3ef2e-618f-11e5-8ac5-06603eb7f303 1.74e-01
Beta5 beta6 beta7 and beta8 integrin cell surface interactions Homo sapiens 108637d6-618e-11e5-8ac5-06603eb7f303 2.09e-01
C-MYC pathway Homo sapiens e3162de9-618e-11e5-8ac5-06603eb7f303 2.72e-01
−log10(p-value)
12hr
NCI-Nature 2016
Regulation of CDC42 activity Homo sapiens 0ffe6681-6195-11e5-8ac5-06603eb7f303 7.70e-03
Regulation of RAC1 activity Homo sapiens 351aacd6-6195-11e5-8ac5-06603eb7f303 1.48e-02
EPHA2 forward signaling Homo sapiens fb172ba9-6191-11e5-8ac5-06603eb7f303 2.44e-02
Sphingosine 1-phosphate (S1P) pathway Homo sapiens eff796f3-6195-11e5-8ac5-06603eb7f303 3.26e-02
Caspase Cascade in Apoptosis Homo sapiens b9d3ef2e-618f-11e5-8ac5-06603eb7f303 3.89e-02
S1P2 pathway Homo sapiens 7796a240-6195-11e5-8ac5-06603eb7f303 4.18e-02
Notch signaling pathway Homo sapiens 88f83518-6194-11e5-8ac5-06603eb7f303 4.63e-02
Reelin signaling pathway Homo sapiens 054f432f-6195-11e5-8ac5-06603eb7f303 5.88e-02
Regulation of p38-alpha and p38-beta Homo sapiens 3062ca15-6195-11e5-8ac5-06603eb7f303 6.25e-02
Aurora A signaling Homo sapiens f131cf8e-618b-11e5-8ac5-06603eb7f303 6.62e-02
−log10(p-value)
24hr
NCI-Nature 2016
IL12 signaling mediated by STAT4 Homo sapiens 72cf19b8-6193-11e5-8ac5-06603eb7f303 1.03e-02
Validated nuclear estrogen receptor alpha network Homo sapiens 58949883-6196-11e5-8ac5-06603eb7f303 1.28e-02
JNK signaling in the CD4+ TCR pathway Homo sapiens 400ebdab-6194-11e5-8ac5-06603eb7f303 1.62e-02
IFN-gamma pathway Homo sapiens 51b1ed75-6193-11e5-8ac5-06603eb7f303 1.88e-02
EPHA2 forward signaling Homo sapiens fb172ba9-6191-11e5-8ac5-06603eb7f303 2.63e-02
IL6-mediated signaling events Homo sapiens e684d5d2-6193-11e5-8ac5-06603eb7f303 2.72e-02
Angiopoietin receptor Tie2-mediated signaling Homo sapiens ad60647c-6188-11e5-8ac5-06603eb7f303 3.04e-02
Signaling events mediated by Stem cell factor receptor (c-Kit) Homo sapiens c6b6861c-6195-11e5-8ac5-06603eb7f303 3.73e-02
RAC1 signaling pathway Homo sapiens faffa4fc-6194-11e5-8ac5-06603eb7f303 4.1e-02
IL2-mediated signaling events Homo sapiens a2a1883c-6193-11e5-8ac5-06603eb7f303 4.29e-02
−log10(p-value)
48hr
NCI-Nature 2016
HIF-1-alpha transcription factor network Homo sapiens 20ef2b81-6193-11e5-8ac5-06603eb7f303 1.41e-02
Regulation of Telomerase Homo sapiens 4dfe97ca-6195-11e5-8ac5-06603eb7f303 1.48e-02
IL6-mediated signaling events Homo sapiens e684d5d2-6193-11e5-8ac5-06603eb7f303 2.7e-02
Signaling events mediated by Stem cell factor receptor (c-Kit) Homo sapiens c6b6861c-6195-11e5-8ac5-06603eb7f303 3.69e-02
SHP2 signaling Homo sapiens 85755aa4-6195-11e5-8ac5-06603eb7f303 4.65e-02
Coregulation of Androgen receptor activity Homo sapiens 27e0e369-6191-11e5-8ac5-06603eb7f303 5.06e-02
Regulation of CDC42 activity Homo sapiens 0ffe6681-6195-11e5-8ac5-06603eb7f303 6.66e-02
EPO signaling pathway Homo sapiens 20fe3c0e-6192-11e5-8ac5-06603eb7f303 7.86e-02
Alternative NF-kappaB pathway Homo sapiens 557add8c-6188-11e5-8ac5-06603eb7f303 8.17e-02
EGF receptor (ErbB1) signaling pathway Homo sapiens NULL 8.28e-02
−log10(p-value)

**Supp Figure 2. Gene ontology for promoters with altered positioning and fuzziness after LPS stimulation**

A. GO for genes with altered nucleosome positioning at each time point following LPS stimulation. Adjusted P values are shown beside ontology terms. Bars in blue indicate significant enrichments. B. GO for genes with altered nucleosome fuzziness at each time point following LPS stimulation.

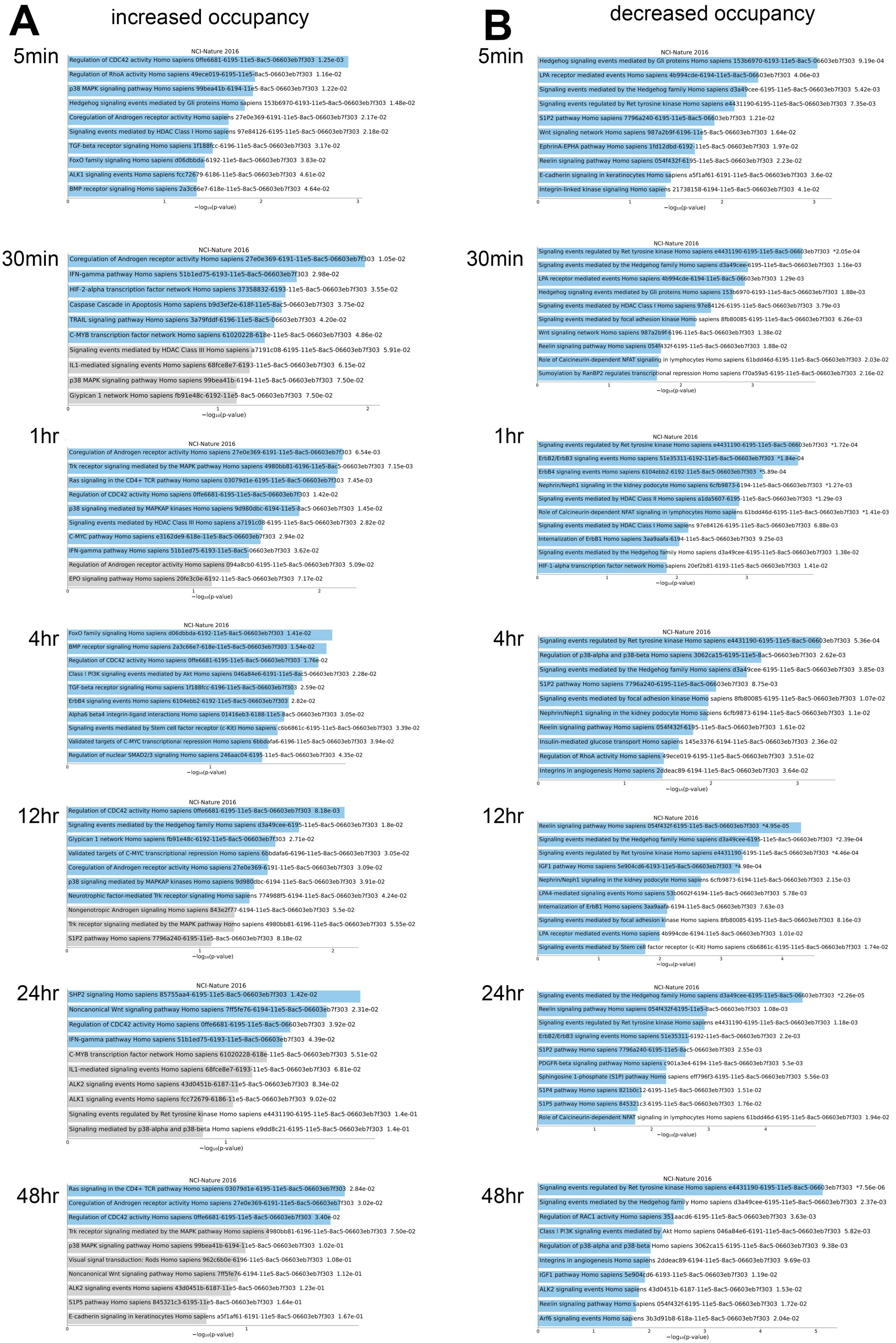

A
increased occupancy
B
decreased occupancy
5min
NCI-Nature 2016
Regulation of CDC42 activity Homo sapiens 0ffe6681-6195-11e5-8ac5-06603eb7f303 1.25e-03
Regulation of RhoA activity Homo sapiens 49ece019-6195-11e5-8ac5-06603eb7f303 1.16e-02
p38 MAPK signaling pathway Homo sapiens 99bea41b-6194-11e5-8ac5-06603eb7f303 1.22e-02
Hedgehog signaling events mediated by Gli proteins Homo sapiens 153b6970-6193-11e5-8ac5-06603eb7f303 1.48e-02
Coregulation of Androgen receptor activity Homo sapiens 27e0e369-6191-11e5-8ac5-06603eb7f303 2.17e-02
Signaling events mediated by HDAC Class I Homo sapiens 97e84126-6195-11e5-8ac5-06603eb7f303 2.18e-02
TGF-beta receptor signaling Homo sapiens 1f188fcc-6196-11e5-8ac5-06603eb7f303 3.17e-02
FoxO family signaling Homo sapiens d06dbbda-6192-11e5-8ac5-06603eb7f303 3.83e-02
ALK1 signaling events Homo sapiens fcc72679-6186-11e5-8ac5-06603eb7f303 4.61e-02
BMP receptor signaling Homo sapiens 2a3c66e7-618e-11e5-8ac5-06603eb7f303 4.64e-02
−log10(p-value)
5min
NCI-Nature 2016
Hedgehog signaling events mediated by Gli proteins Homo sapiens 153b6970-6193-11e5-8ac5-06603eb7f303 9.19e-04
LPA receptor mediated events Homo sapiens 4b994cde-6194-11e5-8ac5-06603eb7f303 4.06e-03
Signaling events mediated by the Hedgehog family Homo sapiens d3a49cee-6195-11e5-8ac5-06603eb7f303 5.42e-03
Signaling events regulated by Ret tyrosine kinase Homo sapiens e4431190-6195-11e5-8ac5-06603eb7f303 7.35e-03
S1P2 pathway Homo sapiens 7796a240-6195-11e5-8ac5-06603eb7f303 1.21e-02
Wnt signaling network Homo sapiens 987a2b9f-6196-11e5-8ac5-06603eb7f303 1.64e-02
EphrinA-EPHA pathway Homo sapiens 1fd12dbd-6192-11e5-8ac5-06603eb7f303 1.97e-02
Reelin signaling pathway Homo sapiens 054f432f-6195-11e5-8ac5-06603eb7f303 2.23e-02
E-cadherin signaling in keratinocytes Homo sapiens a5f1af61-6191-11e5-8ac5-06603eb7f303 3.6e-02
Integrin-linked kinase signaling Homo sapiens 21738158-6194-11e5-8ac5-06603eb7f303 4.1e-02
−log10(p-value)
30min
NCI-Nature 2016
Coregulation of Androgen receptor activity Homo sapiens 27e0e369-6191-11e5-8ac5-06603eb7f303 1.05e-02
IFN-gamma pathway Homo sapiens 51b1ed75-6193-11e5-8ac5-06603eb7f303 2.98e-02
HIF-2-alpha transcription factor network Homo sapiens 37358832-6193-11e5-8ac5-06603eb7f303 3.55e-02
Caspase Cascade in Apoptosis Homo sapiens b9d3ef2e-618f-11e5-8ac5-06603eb7f303 3.75e-02
TRAIL signaling pathway Homo sapiens 3a79fddf-6196-11e5-8ac5-06603eb7f303 4.20e-02
C-MYB transcription factor network Homo sapiens 61020228-618e-11e5-8ac5-06603eb7f303 4.86e-02
Signaling events mediated by HDAC Class III Homo sapiens a7191c08-6195-11e5-8ac5-06603eb7f303 5.91e-02
IL1-mediated signaling events Homo sapiens 68fce8e7-6193-11e5-8ac5-06603eb7f303 6.15e-02
p38 MAPK signaling pathway Homo sapiens 99bea41b-6194-11e5-8ac5-06603eb7f303 7.50e-02
Glypican 1 network Homo sapiens fb91e48c-6192-11e5-8ac5-06603eb7f303 7.50e-02
−log10(p-value)
30min
NCI-Nature 2016
Signaling events regulated by Ret tyrosine kinase Homo sapiens e4431190-6195-11e5-8ac5-06603eb7f303 *2.05e-04
Signaling events mediated by the Hedgehog family Homo sapiens d3a49cee-6195-11e5-8ac5-06603eb7f303 1.16e-03
LPA receptor mediated events Homo sapiens 4b994cde-6194-11e5-8ac5-06603eb7f303 1.29e-03
Hedgehog signaling events mediated by Gli proteins Homo sapiens 153b6970-6193-11e5-8ac5-06603eb7f303 1.88e-03
Signaling events mediated by HDAC Class I Homo sapiens 97e84126-6195-11e5-8ac5-06603eb7f303 3.79e-03
Signaling events mediated by focal adhesion kinase Homo sapiens 8fb80085-6195-11e5-8ac5-06603eb7f303 6.26e-03
Wnt signaling network Homo sapiens 987a2b9f-6196-11e5-8ac5-06603eb7f303 1.38e-02
Reelin signaling pathway Homo sapiens 054f432f-6195-11e5-8ac5-06603eb7f303 1.88e-02
Role of Calcineurin-dependent NFAT signaling in lymphocytes Homo sapiens 61bdd46d-6195-11e5-8ac5-06603eb7f303 2.03e-02
Sumoylation by RanBP2 regulates transcriptional repression Homo sapiens f70a59a5-6195-11e5-8ac5-06603eb7f303 2.16e-02
−log10(p-value)
1hr
NCI-Nature 2016
Coregulation of Androgen receptor activity Homo sapiens 27e0e369-6191-11e5-8ac5-06603eb7f303 6.54e-03
Trk receptor signaling mediated by the MAPK pathway Homo sapiens 4980bb81-6196-11e5-8ac5-06603eb7f303 7.15e-03
Ras signaling in the CD4+ TCR pathway Homo sapiens 03079d1e-6195-11e5-8ac5-06603eb7f303 7.45e-03
Regulation of CDC42 activity Homo sapiens 0ffe6681-6195-11e5-8ac5-06603eb7f303 1.42e-02
p38 signaling mediated by MAPKAP kinases Homo sapiens 9d980dbc-6194-11e5-8ac5-06603eb7f303 1.45e-02
Signaling events mediated by HDAC Class III Homo sapiens a7191c08-6195-11e5-8ac5-06603eb7f303 2.82e-02
C-MYC pathway Homo sapiens e3162de9-618e-11e5-8ac5-06603eb7f303 2.94e-02
IFN-gamma pathway Homo sapiens 51b1ed75-6193-11e5-8ac5-06603eb7f303 3.62e-02
Regulation of Androgen receptor activity Homo sapiens 094a8cb0-6195-11e5-8ac5-06603eb7f303 5.09e-02
EPO signaling pathway Homo sapiens 20fe3c0e-6192-11e5-8ac5-06603eb7f303 7.17e-02
−log10(p-value)
1hr
NCI-Nature 2016
Signaling events regulated by Ret tyrosine kinase Homo sapiens e4431190-6195-11e5-8ac5-06603eb7f303 *1.72e-04
ErbB2/ErbB3 signaling events Homo sapiens 51e35311-6192-11e5-8ac5-06603eb7f303 *1.84e-04
ErbB4 signaling events Homo sapiens 6104ebb2-6192-11e5-8ac5-06603eb7f303 *5.89e-04
Nephrin/Neph1 signaling in the kidney podocyte Homo sapiens 6cfb9873-6194-11e5-8ac5-06603eb7f303 *1.27e-03
Signaling events mediated by HDAC Class II Homo sapiens a1da5607-6195-11e5-8ac5-06603eb7f303 *1.29e-03
Role of Calcineurin-dependent NFAT signaling in lymphocytes Homo sapiens 61bdd46d-6195-11e5-8ac5-06603eb7f303 *1.41e-03
Signaling events mediated by HDAC Class I Homo sapiens 97e84126-6195-11e5-8ac5-06603eb7f303 6.88e-03
Internalization of ErbB1 Homo sapiens 3aa9aafa-6194-11e5-8ac5-06603eb7f303 9.25e-03
Signaling events mediated by the Hedgehog family Homo sapiens d3a49cee-6195-11e5-8ac5-06603eb7f303 1.38e-02
HIF-1-alpha transcription factor network Homo sapiens 20ef2b81-6193-11e5-8ac5-06603eb7f303 1.41e-02
−log10(p-value)
4hr
NCI-Nature 2016
FoxO family signaling Homo sapiens d06dbbda-6192-11e5-8ac5-06603eb7f303 1.41e-02
BMP receptor signaling Homo sapiens 2a3c66e7-618e-11e5-8ac5-06603eb7f303 1.54e-02
Regulation of CDC42 activity Homo sapiens 0ffe6681-6195-11e5-8ac5-06603eb7f303 1.76e-02
Class I PI3K signaling events mediated by Akt Homo sapiens 046a84e6-6191-11e5-8ac5-06603eb7f303 2.28e-02
TGF-beta receptor signaling Homo sapiens 1f188fcc-6196-11e5-8ac5-06603eb7f303 2.59e-02
ErbB4 signaling events Homo sapiens 6104ebb2-6192-11e5-8ac5-06603eb7f303 2.82e-02
Alpha6 beta4 integrin-ligand interactions Homo sapiens 01416eb3-6188-11e5-8ac5-06603eb7f303 3.05e-02
Signaling events mediated by Stem cell factor receptor (c-Kit) Homo sapiens c6b6861c-6195-11e5-8ac5-06603eb7f303 3.39e-02
Validated targets of C-MYC transcriptional repression Homo sapiens 6bbdafa6-6196-11e5-8ac5-06603eb7f303 3.94e-02
Regulation of nuclear SMAD2/3 signaling Homo sapiens 246aac04-6195-11e5-8ac5-06603eb7f303 4.35e-02
−log10(p-value)
4hr
NCI-Nature 2016
Signaling events regulated by Ret tyrosine kinase Homo sapiens e4431190-6195-11e5-8ac5-06603eb7f303 5.36e-04
Regulation of p38-alpha and p38-beta Homo sapiens 3062ca15-6195-11e5-8ac5-06603eb7f303 2.62e-03
Signaling events mediated by the Hedgehog family Homo sapiens d3a49cee-6195-11e5-8ac5-06603eb7f303 3.85e-03
S1P2 pathway Homo sapiens 7796a240-6195-11e5-8ac5-06603eb7f303 8.75e-03
Signaling events mediated by focal adhesion kinase Homo sapiens 8fb80085-6195-11e5-8ac5-06603eb7f303 1.07e-02
Nephrin/Neph1 signaling in the kidney podocyte Homo sapiens 6cfb9873-6194-11e5-8ac5-06603eb7f303 1.1e-02
Reelin signaling pathway Homo sapiens 054f432f-6195-11e5-8ac5-06603eb7f303 1.61e-02
Insulin-mediated glucose transport Homo sapiens 145e3376-6194-11e5-8ac5-06603eb7f303 2.36e-02
Regulation of RhoA activity Homo sapiens 49ece019-6195-11e5-8ac5-06603eb7f303 3.51e-02
Integrins in angiogenesis Homo sapiens 2ddeac89-6194-11e5-8ac5-06603eb7f303 3.64e-02
−log10(p-value)
12hr
NCI-Nature 2016
Regulation of CDC42 activity Homo sapiens 0ffe6681-6195-11e5-8ac5-06603eb7f303 8.18e-03
Signaling events mediated by the Hedgehog family Homo sapiens d3a49cee-6195-11e5-8ac5-06603eb7f303 1.8e-02
Glypican 1 network Homo sapiens fb91e48c-6192-11e5-8ac5-06603eb7f303 2.71e-02
Validated targets of C-MYC transcriptional repression Homo sapiens 6bbdafa6-6196-11e5-8ac5-06603eb7f303 3.05e-02
Coregulation of Androgen receptor activity Homo sapiens 27e0e369-6191-11e5-8ac5-06603eb7f303 3.09e-02
p38 signaling mediated by MAPKAP kinases Homo sapiens 9d980dbc-6194-11e5-8ac5-06603eb7f303 3.91e-02
Neurotrophic factor-mediated Trk receptor signaling Homo sapiens 774988f5-6194-11e5-8ac5-06603eb7f303 4.24e-02
Nongenotropic Androgen signaling Homo sapiens 843e2f77-6194-11e5-8ac5-06603eb7f303 5.5e-02
Trk receptor signaling mediated by the MAPK pathway Homo sapiens 4980bb81-6196-11e5-8ac5-06603eb7f303 5.55e-02
S1P2 pathway Homo sapiens 7796a240-6195-11e5-8ac5-06603eb7f303 8.18e-02
−log10(p-value)
12hr
NCI-Nature 2016
Reelin signaling pathway Homo sapiens 054f432f-6195-11e5-8ac5-06603eb7f303 *4.95e-05
Signaling events mediated by the Hedgehog family Homo sapiens d3a49cee-6195-11e5-8ac5-06603eb7f303 *2.39e-04
Signaling events regulated by Ret tyrosine kinase Homo sapiens e4431190-6195-11e5-8ac5-06603eb7f303 *4.46e-04
IGF1 pathway Homo sapiens 5e904cd6-6193-11e5-8ac5-06603eb7f303 *4.98e-04
Nephrin/Neph1 signaling in the kidney podocyte Homo sapiens 6cfb9873-6194-11e5-8ac5-06603eb7f303 2.15e-03
LPA4-mediated signaling events Homo sapiens 53b0602f-6194-11e5-8ac5-06603eb7f303 5.78e-03
Internalization of ErbB1 Homo sapiens 3aa9aafa-6194-11e5-8ac5-06603eb7f303 7.63e-03
Signaling events mediated by focal adhesion kinase Homo sapiens 8fb80085-6195-11e5-8ac5-06603eb7f303 8.16e-03
LPA receptor mediated events Homo sapiens 4b994cde-6194-11e5-8ac5-06603eb7f303 1.01e-02
Signaling events mediated by Stem cell factor receptor (c-Kit) Homo sapiens c6b6861c-6195-11e5-8ac5-06603eb7f303 1.74e-02
−log10(p-value)
24hr
NCI-Nature 2016
SHP2 signaling Homo sapiens 85755aa4-6195-11e5-8ac5-06603eb7f303 1.42e-02
Noncanonical Wnt signaling pathway Homo sapiens 7ff5fe76-6194-11e5-8ac5-06603eb7f303 2.31e-02
Regulation of CDC42 activity Homo sapiens 0ffe6681-6195-11e5-8ac5-06603eb7f303 3.92e-02
IFN-gamma pathway Homo sapiens 51b1ed75-6193-11e5-8ac5-06603eb7f303 4.39e-02
C-MYB transcription factor network Homo sapiens 61020228-618e-11e5-8ac5-06603eb7f303 5.51e-02
IL1-mediated signaling events Homo sapiens 68fce8e7-6193-11e5-8ac5-06603eb7f303 6.81e-02
ALK2 signaling events Homo sapiens 43d0451b-6187-11e5-8ac5-06603eb7f303 8.34e-02
ALK1 signaling events Homo sapiens fcc72679-6186-11e5-8ac5-06603eb7f303 9.02e-02
Signaling events regulated by Ret tyrosine kinase Homo sapiens e4431190-6195-11e5-8ac5-06603eb7f303 1.4e-01
Signaling mediated by p38-alpha and p38-beta Homo sapiens e9dd8c21-6195-11e5-8ac5-06603eb7f303 1.4e-01
−log10(p-value)
24hr
NCI-Nature 2016
Signaling events mediated by the Hedgehog family Homo sapiens d3a49cee-6195-11e5-8ac5-06603eb7f303 *2.26e-05
Reelin signaling pathway Homo sapiens 054f432f-6195-11e5-8ac5-06603eb7f303 1.08e-03
Signaling events regulated by Ret tyrosine kinase Homo sapiens e4431190-6195-11e5-8ac5-06603eb7f303 1.18e-03
ErbB2/ErbB3 signaling events Homo sapiens 51e35311-6192-11e5-8ac5-06603eb7f303 2.2e-03
S1P2 pathway Homo sapiens 7796a240-6195-11e5-8ac5-06603eb7f303 2.55e-03
PDGFR-beta signaling pathway Homo sapiens c901a3e4-6194-11e5-8ac5-06603eb7f303 5.5e-03
Sphingosine 1-phosphate (S1P) pathway Homo sapiens eff796f3-6195-11e5-8ac5-06603eb7f303 5.56e-03
S1P4 pathway Homo sapiens 821b0c12-6195-11e5-8ac5-06603eb7f303 1.51e-02
S1P5 pathway Homo sapiens 845321c3-6195-11e5-8ac5-06603eb7f303 1.76e-02
Role of Calcineurin-dependent NFAT signaling in lymphocytes Homo sapiens 61bdd46d-6195-11e5-8ac5-06603eb7f303 1.94e-02
−log10(p-value)
48hr
NCI-Nature 2016
Ras signaling in the CD4+ TCR pathway Homo sapiens 03079d1e-6195-11e5-8ac5-06603eb7f303 2.84e-02
Coregulation of Androgen receptor activity Homo sapiens 27e0e369-6191-11e5-8ac5-06603eb7f303 3.02e-02
Regulation of CDC42 activity Homo sapiens 0ffe6681-6195-11e5-8ac5-06603eb7f303 3.40e-02
Trk receptor signaling mediated by the MAPK pathway Homo sapiens 4980bb81-6196-11e5-8ac5-06603eb7f303 7.50e-02
p38 MAPK signaling pathway Homo sapiens 99bea41b-6194-11e5-8ac5-06603eb7f303 1.02e-01
Visual signal transduction: Rods Homo sapiens 962c6b0e-6196-11e5-8ac5-06603eb7f303 1.08e-01
Noncanonical Wnt signaling pathway Homo sapiens 7ff5fe76-6194-11e5-8ac5-06603eb7f303 1.12e-01
ALK2 signaling events Homo sapiens 43d0451b-6187-11e5-8ac5-06603eb7f303 1.23e-01
S1P5 pathway Homo sapiens 845321c3-6195-11e5-8ac5-06603eb7f303 1.64e-01
E-cadherin signaling in keratinocytes Homo sapiens a5f1af61-6191-11e5-8ac5-06603eb7f303 1.67e-01
−log10(p-value)
48hr
NCI-Nature 2016
Signaling events regulated by Ret tyrosine kinase Homo sapiens e4431190-6195-11e5-8ac5-06603eb7f303 *7.56e-06
Signaling events mediated by the Hedgehog family Homo sapiens d3a49cee-6195-11e5-8ac5-06603eb7f303 2.37e-03
Regulation of RAC1 activity Homo sapiens 351aacd6-6195-11e5-8ac5-06603eb7f303 3.63e-03
Class I PI3K signaling events mediated by Akt Homo sapiens 046a84e6-6191-11e5-8ac5-06603eb7f303 5.82e-03
Regulation of p38-alpha and p38-beta Homo sapiens 3062ca15-6195-11e5-8ac5-06603eb7f303 9.38e-03
Integrins in angiogenesis Homo sapiens 2ddeac89-6194-11e5-8ac5-06603eb7f303 9.69e-03
IGF1 pathway Homo sapiens 5e904cd6-6193-11e5-8ac5-06603eb7f303 1.19e-02
ALK2 signaling events Homo sapiens 43d0451b-6187-11e5-8ac5-06603eb7f303 1.53e-02
Reelin signaling pathway Homo sapiens 054f432f-6195-11e5-8ac5-06603eb7f303 1.72e-02
Arf6 signaling events Homo sapiens 3b3d91b8-618a-11e5-8ac5-06603eb7f303 2.04e-02
−log10(p-value)

**Supp Figure 3. Gene ontology for promoters with altered nucleosome occupancy after LPS stimulation**

A. GO for genes with increased nucleosome occupancy at each time point following LPS stimulation. Adjusted P values are shown beside ontology terms. Bars in blue indicate significant enrichments. B. GO for genes with decreased nucleosome occupancy at each time point following LPS stimulation.

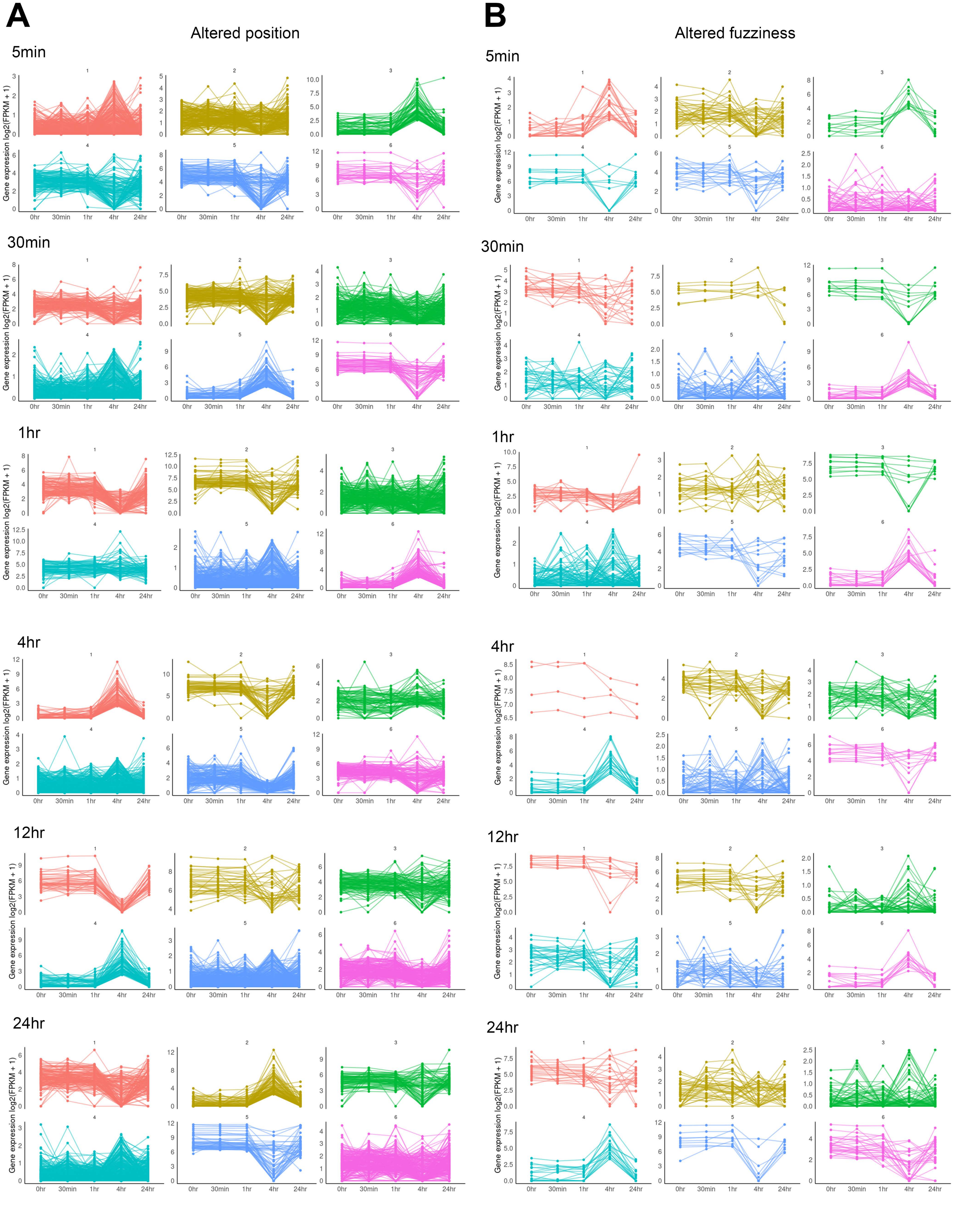
A
Altered position
B
Altered fuzziness
5min
30min
1hr
4hr
12hr
24hr
Gene expression log2(FPKM + 1)
0hr
30min
1hr
4hr
24hr

**Supp. Figure 4. Gene expression patterns for genes with altered nucleosome positioning and fuzziness**

A. Kmeans clustered line plots of gene expression (Log2(FPKM+1)) for promoters with altered nucleosome distribution patterns at each time point following LPS stimulation. B. Kmeans clustered line plots of gene expression (Log2(FPKM+1)) for promoters with altered nucleosome fuzziness patterns at each time point following LPS stimulation.

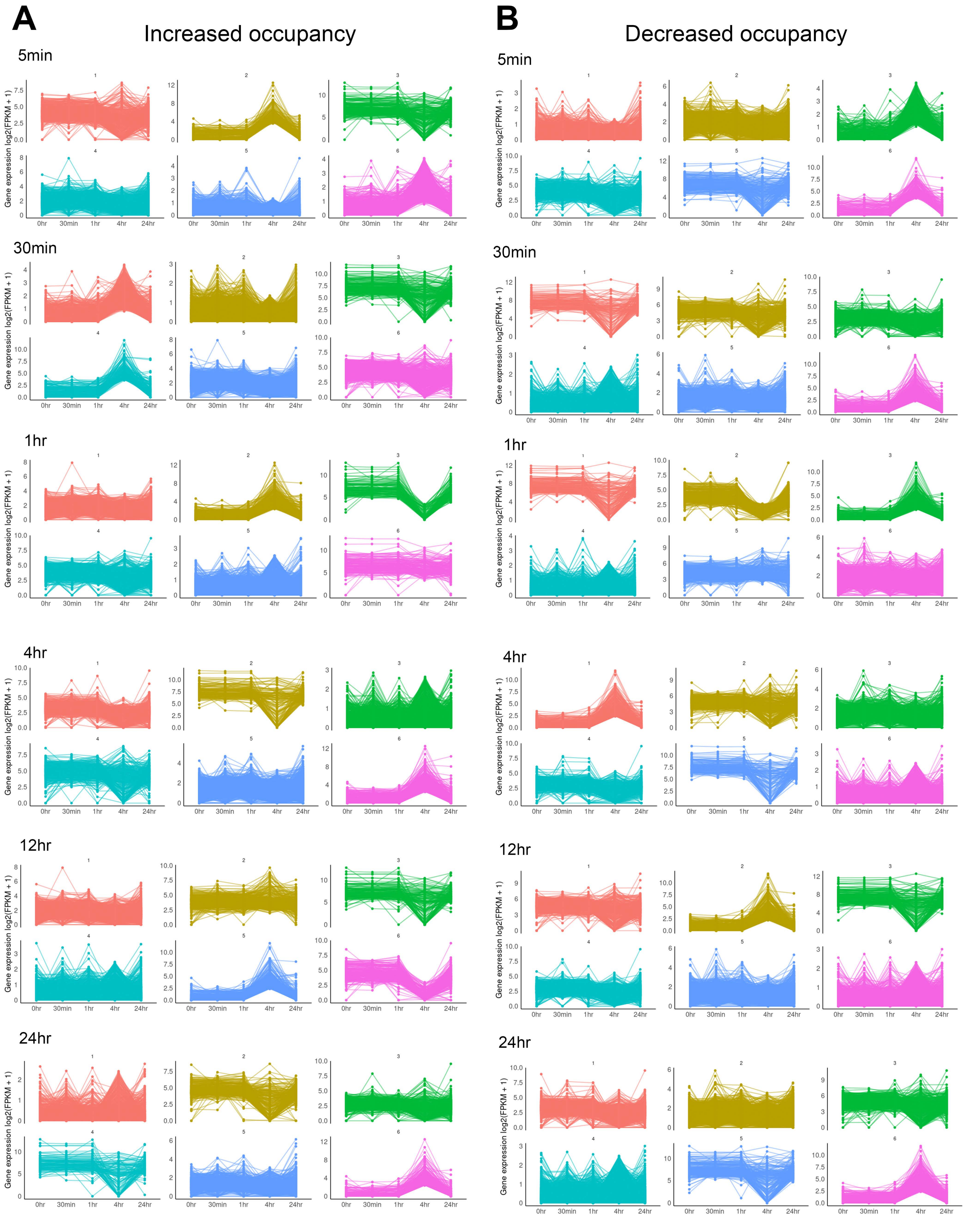
A
Increased occupancy
B
Decreased occupancy
5min
30min
1hr
4hr
12hr
24hr
Gene expression log2(FPKM + 1)
0hr
30min
1hr
4hr
24hr

**Supp. Figure 5. Gene expression patterns for genes with altered nucleosome occupancy**

A. Kmeans clustered line plots of gene expression (Log2(FPKM+1)) for promoters with increased nucleosome occupancy patterns at each time point following LPS stimulation. B. Kmeans clustered line plots of gene expression (Log2(FPKM+1)) for promoters with decreased nucleosome occupancy patterns at each time point following LPS stimulation.

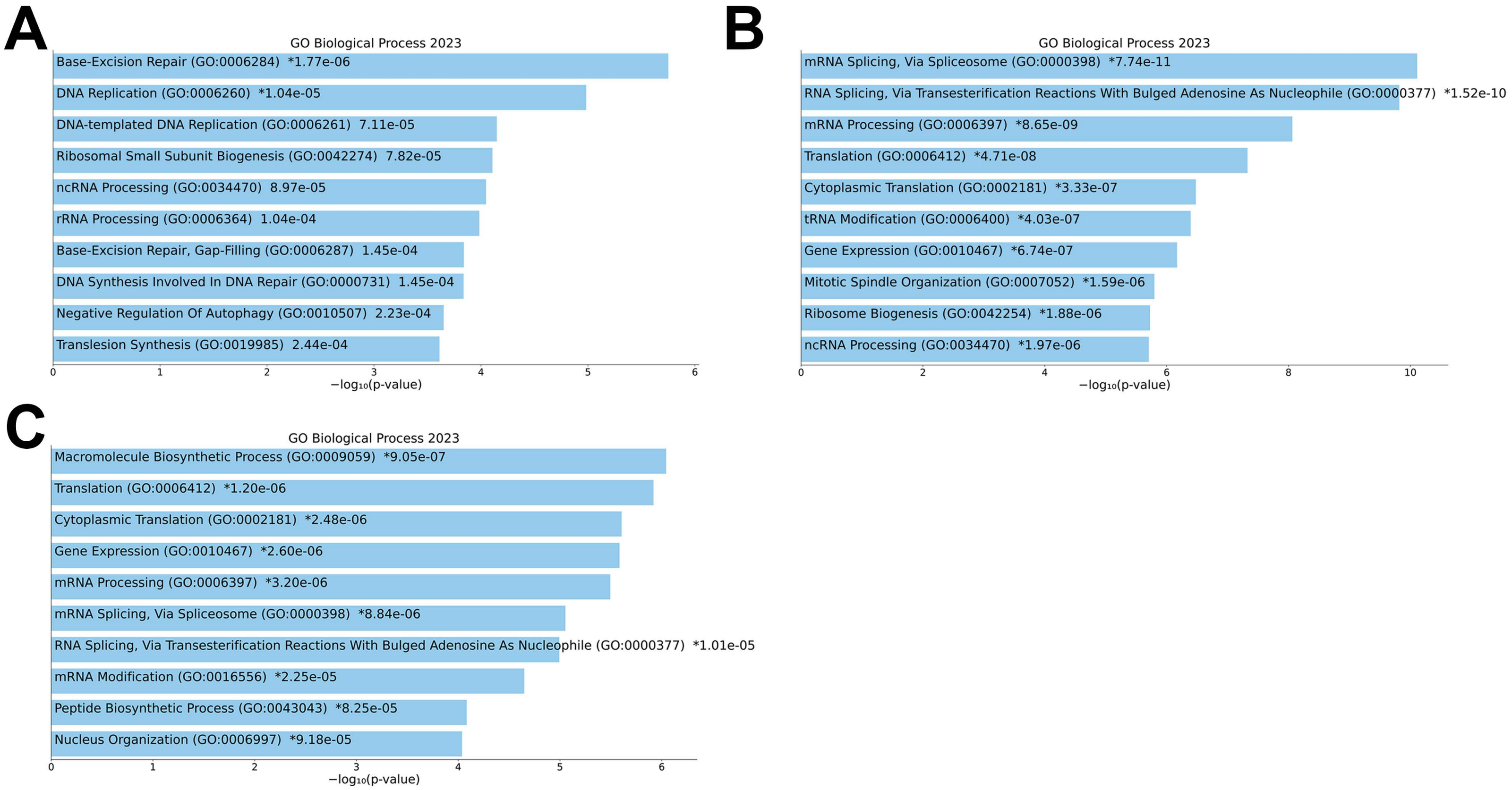
A
GO Biological Process 2023
Base-Excision Repair (GO:0006284) *1.77e-06
DNA Replication (GO:0006260) *1.04e-05
DNA-templated DNA Replication (GO:0006261) 7.11e-05
Ribosomal Small Subunit Biogenesis (GO:0042274) 7.82e-05
ncRNA Processing (GO:0034470) 8.97e-05
rRNA Processing (GO:0006364) 1.04e-04
Base-Excision Repair, Gap-Filling (GO:0006287) 1.45e-04
DNA Synthesis Involved In DNA Repair (GO:0000731) 1.45e-04
Negative Regulation Of Autophagy (GO:0010507) 2.23e-04
Translesion Synthesis (GO:0019985) 2.44e-04
−log₁₀(p-value)
B
GO Biological Process 2023
mRNA Splicing, Via Spliceosome (GO:0000398) *7.74e-11
RNA Splicing, Via Transesterification Reactions With Bulged Adenosine As Nucleophile (GO:0000377) *1.52e-10
mRNA Processing (GO:0006397) *8.65e-09
Translation (GO:0006412) *4.71e-08
Cytoplasmic Translation (GO:0002181) *3.33e-07
tRNA Modification (GO:0006400) *4.03e-07
Gene Expression (GO:0010467) *6.74e-07
Mitotic Spindle Organization (GO:0007052) *1.59e-06
Ribosome Biogenesis (GO:0042254) *1.88e-06
ncRNA Processing (GO:0034470) *1.97e-06
−log₁₀(p-value)
C
GO Biological Process 2023
Macromolecule Biosynthetic Process (GO:0009059) *9.05e-07
Translation (GO:0006412) *1.20e-06
Cytoplasmic Translation (GO:0002181) *2.48e-06
Gene Expression (GO:0010467) *2.60e-06
mRNA Processing (GO:0006397) *3.20e-06
mRNA Splicing, Via Spliceosome (GO:0000398) *8.84e-06
RNA Splicing, Via Transesterification Reactions With Bulged Adenosine As Nucleophile (GO:0000377) *1.01e-05
mRNA Modification (GO:0016556) *2.25e-05
Peptide Biosynthetic Process (GO:0043043) *8.25e-05
Nucleus Organization (GO:0006997) *9.18e-05
−log₁₀(p-value)

**Supp. Figure 6. Gene ontology of highly positioned sensitive clusters**

A. Biological Process gene ontology with adjusted p values for common genes in the positioned sensitive clusters at 24hr and 48hr. B. Biological Process gene ontology with adjusted p values for genes in the positioned sensitivity clusters that were unique to 24hr. C. Biological Process gene ontology with adjusted p values for genes in the positioned sensitivity clusters that were unique to 48hr.

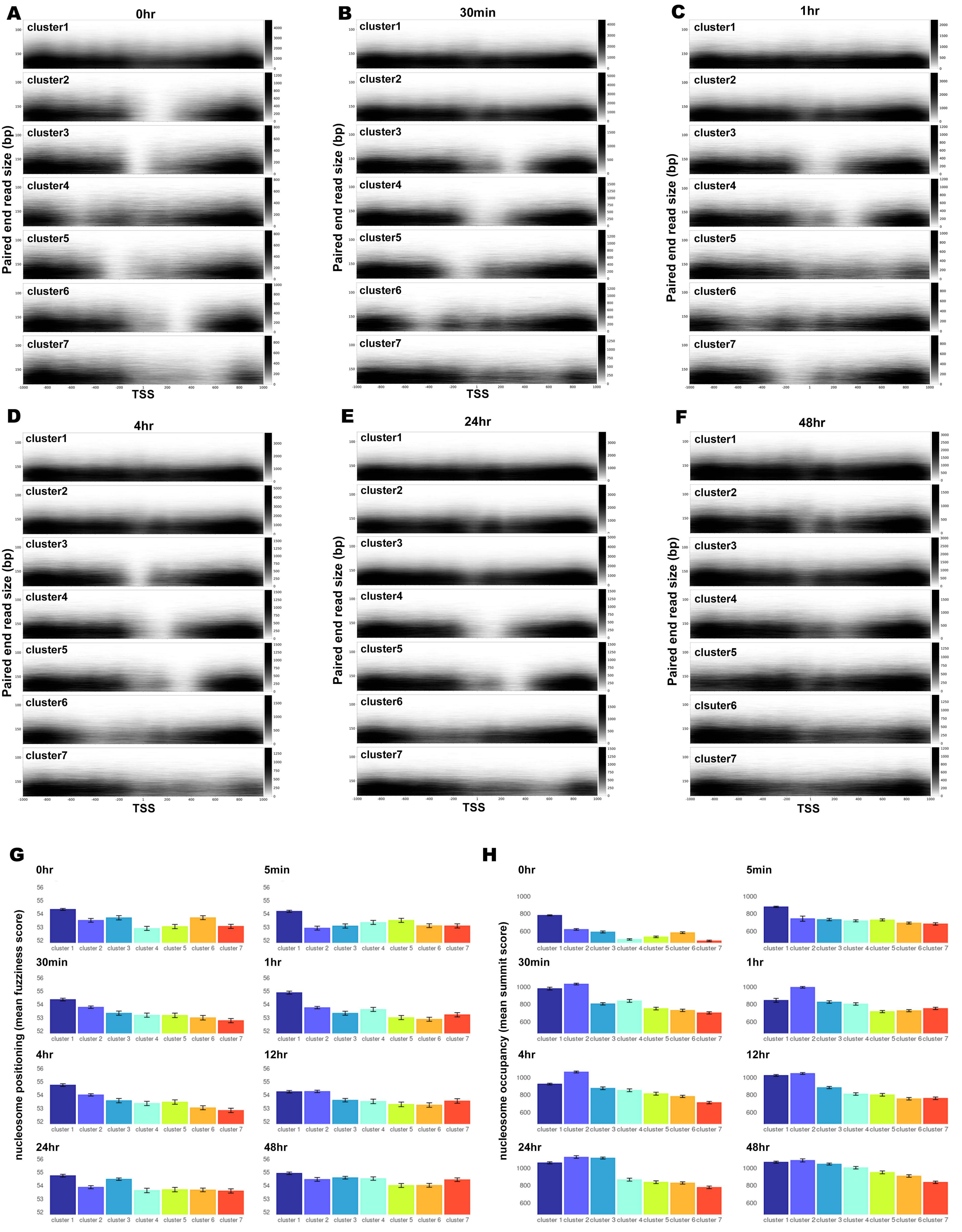

A
0hr
B
30min
C
1hr
D
4hr
E
24hr
F
48hr
cluster1
cluster2
cluster3
cluster4
cluster5
cluster6
clsuter6
cluster7
Paired end read size (bp)
TSS
-1000
-800
-600
-400
-200
1
200
400
600
800
1000
G
0hr
5min
30min
1hr
4hr
12hr
24hr
48hr
nucleosome positioning (mean fuzziness score)
cluster 1
cluster 2
cluster 3
cluster 4
cluster 5
cluster 6
cluster 7
H
nucleosome occupancy (mean summit score)

**Supp. Figure 7 Characteristics of sensitive nucleosomes**

A. Fragmaps show the distribution of paired end fragments from light digests by size over a 2kb region centered on the TSS of promoters for each sensitivity cluster at 0hr. B. The same for 30 min after LPS stimulation. C. The same for 1hr after LPS stimulation. D. The same for 4hr after LPS stimulation. E. The same for 24hr after LPS stimulation. F. The same for 48hr after LPS stimulation. G. Average nucleosome occupancy, as measured by fuzzines scores, with error bars showing 95% confidence intervals for sensitivity clusters following LPS stimulation. H. Average nucleosome positioning scores, as measured by summit scores, for sensitivity clusters following LPS stimulation at all time points with error bars showing 95% confidence intervals.

### Supplemental Tables (see separate excel sheets for tables 1-4)

#### Supp. Table 1

List of genes with one or more nucleosomes within promoter regions with altered positioning at one or more time points following LPS stimulation compared to unstimulated cells. We defined nucleosomes as being repositioned if there was a shift in the nucleosome summit position greater than 80bp between time points.

#### Supp. Table 2

List of genes with one or more nucleosomes within promoter regions with altered fuzziness at one or more time points following LPS stimulation compared to unstimulated cells. We considered a change in nucleosome fuzziness to occur if there was a log2 fold change greater than 1.5 between conditions combined with a p-value and FDR under 0.05.

#### Supp. Table 3

List of genes with one or more nucleosomes within promoter regions with increased occupancy at one or more time points following LPS stimulation compared to unstimulated cells. We defined alterations in nucleosome occupancy as having a log2 fold change greater than 5 with a p-value and FDR under 0.05 for any time point compared to the basal occupancy seen in 0hr.

#### Supp. Table 4

List of genes with one or more nucleosomes within promoter regions with decreased occupancy at one or more time points following LPS stimulation compared to unstimulated cells. We defined alterations in nucleosome occupancy as having a log2 fold change less than 5 with a p-value and FDR under 0.05 for any time point compared to the basal occupancy seen in 0hr.

#### Supp. Table 5.

Percent of genes with repositioned nucleosomes that are differentially or non-differentially expressed over time.

| | Altered position | | Altered fuzziness | | Increased occupancy | | Decreased occupancy | |
|---|---|---|---|---|---|---|---|---|
| time | DEGs | non-DEGs | DEGs | non-DEGs | DEGs | non-DEGs | DEGs | non-DEGs |
| 5min | 29.5 | 70.5 | 32.1 | 67.9 | 30.8 | 69.2 | 31.5 | 68.5 |
| 30min | 28.3 | 71.7 | 29.4 | 70.6 | 31.4 | 68.6 | 31.2 | 68.8 |
| 1hr | 29.0 | 71.0 | 29.7 | 70.3 | 30.7 | 69.3 | 31.3 | 68.7 |
| 4hr | 28.5 | 71.5 | 25.9 | 74.1 | 32.2 | 67.8 | 31.1 | 68.9 |
| 12hr | 27.1 | 72.9 | 31.3 | 68.7 | 30.9 | 69.1 | 32.1 | 67.9 |
| 24hr | 29.1 | 70.9 | 33.0 | 67.0 | 29.8 | 70.2 | 27.5 | 72.5 |
| 48hr | 30.3 | 69.7 | 38.2 | 61.8 | 31.2 | 68.8 | 31.0 | 69.0 |
| all genes | 29.7 | 70.3 | 29.7 | 70.3 | 29.7 | 70.3 | 29.7 | 70.3 |

#### Supp. Table 6.

Percent of genes with one or more statistically significant nucleosomes coded as sensitive, resistant, or neutral that are differentially or non-differentially expressed over time.

| | Sensitive | | Resistant | | Neutral | |
|---|---|---|---|---|---|---|
| time | DEGs | non-DEGs | DEGs | non-DEGs | DEGs | non-DEGs |
| 0min | 28.5 | 71.5 | 30.7 | 69.3 | 29.8 | 70.2 |
| 5min | 28.6 | 71.4 | 31.0 | 69.0 | 29.8 | 70.2 |
| 30min | 28.7 | 71.3 | 31.3 | 68.7 | 29.8 | 70.2 |
| 1hr | 28.1 | 71.9 | 31.2 | 68.8 | 29.7 | 70.3 |
| 4hr | 29.0 | 71.0 | 31.0 | 69.0 | 29.8 | 70.2 |
| 12hr | 29.7 | 70.3 | 30.4 | 69.6 | 29.8 | 70.2 |
| 24hr | 29.3 | 70.7 | 31.0 | 69.0 | 29.8 | 70.2 |
| 48hr | 30.0 | 70.0 | 29.4 | 70.6 | 29.8 | 70.2 |

#### Alt text for figures and tables

Table 1: A table showing the number of genes with nucleosomes that are repositioned, have altered fuzziness, have increased occupancy, or decreased occupancy over each time point compared to unstimulated macrophages. The table includes columns for time, repositioning, altered fuzziness, increased occupancy, and decreased occupancy. The number of genes with repositioned nucleosomes remains relatively stable, with a slight increase at 24 and 48 hours. Altered fuzziness shows minor fluctuations, peaking at 1 hour. Increased occupancy is highest at 4 hours and decreases by 24 and 48 hours. Decreased occupancy is initially high, peaks at 24 hours, and remains elevated at 48 hours.

Table 2: Table showing the percent of genes with repositioned nucleosomes that are differentially expressed (DEGs) or non-differentially expressed (non-DEGs) over time. Overall, the percentage of DEGs (approximately 30%) remains relatively stable across different time points regardless if the promoter has a change in position, fuzziness, or occupancy.

Table 3: Table showing the number and percent of sensitive and resistant nucleosomes within promoters at each time point following LPS stimulation. Key trends include an increase in the percentage of sensitive nucleosomes at 1 hour (16.4%) and 48 hours (15.6%) compared to the initial time point (11.9%). The percentage of resistant nucleosomes decreases over time, with the highest percentage at 0 hours (15.6%) and the lowest at 12 hours (8%). Neutral nucleosomes remain the majority at all time points.

Graphical abstract: A graphical abstract showing an experimental schematic of THP-1 derived macrophages being treated with LPS and harvested at the following time points: 0hr, 5min, 30min, 1hr, 4hr, 12hr, 24hr, and 48hr. A pie chart shows that the majority of promoters have static nucleosome positions after LPS stimulation with approximately seven percent of promoters undergoing a change in position and approximately 20 percent of promoters undergoing a change in occupancy after LPS stimulation. The redistribution of nucleosomes over a subset of response genes is associated with expression of these genes after LPS stimulation. A pie chart shows that over sixty percent of promoters become increasingly sensitive after LPS stimulation with less than ten percent of genes having a static sensitivity profile across the time course. While most genes experience alterations in sensitivity, only a subset of these genes are differentially expressed following LPS stimulation.

Figure 1. Schematic representation of the MNase based methods to probe nucleosome distribution and sensitivity. The experimental time course of treating THP-1 derived macrophages with LPS is shown along with a schematic of crosslinking cells, quenching excess formaldehyde, isolating nuclei, digesting with MNase, size selecting mononucleosomes, and sequencing libraries. A schematic of a promoter is shown with nucleosomes that are either sensitive or resistant to MNase digestion. Sensitive nucleosomes are preferentially released under light digestion while resistant nucleosomes are released under heavy digestion. Both sensitive and resistant nucleosomes are combined to generate nucleosome distribution maps while the ratio of sensitive over resistant nucleosomes is used to determine overall nucleosome sensitivity.

Figure 2. Panel A shows a representative agarose gel of DNA isolated from heavy and light MNase digests of crosslinked chromatin. A nucleosomal ladder is visible in both heavy and light digest conditions with larger fragments at the limit of resolution present in light but not heavy digests. A label indicates 150bp on the molecular weight marker which is the approximate size of mononucleosome fragments on the gel. Panel B shows heatmaps of nucleosome distribution signal over a 2kb region centered on the TSS of pol II genes. Average line plots above each heatmap show visually similar canonical nucleosome distribution patterns at all time points. Panel C shows average nucleosome distribution patterns at all time points overlaid in a single line plot with 95% confidence intervals. Lower occupancy over the +1 nucleosome is apparent in unstimulated cells compared with all other time points. Panel D shows nucleosome distribution heatmaps with seven promoter classes defined by kmeans clustering of the 0hr time point. Overall patterns are similar over time with the exception of increased occupancy of the +1 nucleosome following LPS stimulation. Panel E shows graphical representation of nucleosome distribution alterations including repositioning, altered fuzziness, and increased or decreased occupancy. Panel F shows an upset plot of genes with repositioned nucleosomes. The majority of which are unique to a single time point. Panel G shows an upset plot of genes with altered nucleosome fuzziness, the majority of which are unique to a single time point. Panel H shows an upset plot of genes with increased nucleosome occupancy, many of which are common to multiple time points. Panel I shows an upset plot of genes with decreased nucleosome occupancy many of which are common to multiple time points..

Figure 3. Panel A shows a line plot of nucleosome distribution with 95% confidence intervals over LPS response and matched gene sets over each time point. Panel B shows boxplots of gene expression (log2(FPKM+1)) for LPS response genes after stimulation. Expression increases marginally 1 and 24 hours after stimulation. Panel C shows line plots of average nucleosome distribution with 95% confidence intervals over early response genes as defined by expression profiles and matched gene sets. Panel D shows boxplots of gene expression for early LPS response genes with increased expression 30min and 1he after stimulation. Panel E shows line plots of average nucleosome distribution with 95% confidence intervals over late response genes and matched gene sets. Panel F shows boxplots of gene expression for late LPS response genes which increase expression 4hr and 24hr after stimulation. Panel G shows heatmaps of nucleosome distribution over all promoters grouped by gene expression quartiles at 0hr, 30min, 1h4, 4hr, and 24hr after LPS stimulation. Alluvial diagrams show changes in gene clustering between time points. Panel H shows nucleosome distribution over the HLA-DMA promoter over each time point. Panel I shows nucleosome distribution over the IL-10 promoter over all time points. Panel J shows nucleosome distribution over the SOD2 promoter over all time points. Panel K shows nucleosome distribution over the ISG15 promoter over all time points.

Figure 4. Panel A shows heatmaps of nucleosome sensitivity centered on the TSSs of pol II genes over all time points grouped by kmeans clustering of unstimulated macrophages. Line plots above each plot show average signal values for each cluster. There is a trend towards reduced resistance following LPS stimulation. Panel B shows Pearson correlation scores for nucleosome sensitivity for all time points over promoters calculated in 10bp bins. Correlation coefficients range from 0.12 to 0.33 for all samples. Panel C shows heatmaps of nucleosome sensitivity over all time points with promoters grouped by kmeans clustering of each independent time point. Each heatmap shows several positioned resistant clusters and a diffusely sensitive cluster. Panel D shows nucleosome sensitivity plotted over the promoters of pol II genes separated into gene expression quartiles over 0hr, 30min, 1hr, 4hr, and 24hr samples. Highly expressed genes tend to have a sensitive -1 nucleosome and a resistant +1 nucleosome at all time points. Panel E shows average nucleosome sensitivity profiles as line plots with 95% confidence intervals over primary LPS response and matched set of non-response genes over all time points. The signal of matched and response genes is overlapping at all time points. Panel F shows nucleosome sensitivity profiles over the SOD2 promoter at all time points. Panel G shows nucleosome sensitivity over the ISG15 promoter at all time points.

Figure 5. Panel A shows nucleosome distribution, sensitivity, H3K27ac ChIP, and ATAC signal over promoters in unstimulated macrophages clustered based on nucleosome sensitivity. There is reduced ATAC and H3K27ac signal over the diffusely sensitive cluster. Panel B shows nucleosome distribution, sensitivity, H3K27ac ChIP, and ATAC signal over promoters in macrophages 30 minutes after LPS stimulation clustered based on nucleosome sensitivity. There is reduced ATAC and H3K27ac signal over the diffusely sensitive cluster. Panel C shows nucleosome distribution, sensitivity, H3K27ac ChIP, and ATAC signal over promoters in macrophages 1 hour after LPS stimulation clustered based on nucleosome sensitivity. There is reduced ATAC and H3K27ac signal over the diffusely sensitive cluster. Panel D shows nucleosome distribution, sensitivity, H3K27ac ChIP, and ATAC signal over promoters in macrophages 4 hours after LPS stimulation clustered based on nucleosome sensitivity. There is reduced ATAC and H3K27ac signal over the diffusely sensitive cluster. Panel E shows nucleosome distribution, sensitivity, H3K27ac ChIP, and ATAC signal over promoters in macrophages 24 hours after LPS stimulation clustered based on nucleosome sensitivity. There is reduced ATAC and H3K27ac signal over the diffusely sensitive cluster and greatest ATAC and H3K27ac signal over the highly positioned sensitive cluster. Panel F shows boxplots of gene expression (log2(FPKM+1)) for each sensitivity cluster in unstimulated macrophages. The greatest expression is seen over positioned resistant clusters. Panel G shows boxplots of gene expression for each sensitivity cluster in macrophages 30 minutes after LPS stimulation. The greatest expression is seen over positioned resistant clusters. Panel H shows boxplots of gene expression for each sensitivity cluster in macrophages one hour after LPS stimulation. The greatest expression is seen over positioned resistant clusters. Panel I shows boxplots of gene expression for each sensitivity cluster in macrophages 4 hours after LPS stimulation. The greatest expression is seen over positioned resistant clusters. Panel J shows boxplots of gene expression for each sensitivity cluster in macrophages 24 hours after LPS stimulation. The greatest expression is seen over the positioned sensitive cluster. Panel K shows a heatmap of nucleosome sensitivity z scores across promoters of primary LPS response genes. The majority of promoters are initially resistant but become more sensitive over time. Panel L shows a heatmap of nucleosome occupancy z scores over the same response genes with mixed signals over time. Panel M shows a heatmap of gene expression z scores for the same set of genes with a trend of increasing expression over time.

Figure 6. Panel A shows nucleosome distribution, nucleosome sensitivity, ATAC signal, and H3K27ac signal over IL-1B, ICAM1, NFKBIA, and IFIT3 promoters. Signal for each promoter is shown over a 2kb region centered on the TSS and signal values for all datatypes have fixed axes at all time points. There is a general trend of nucleosome redistribution, increased sensitivity, and increases in both ATACseq and H3K27ac signal over time. Panel B shows a heatmap of gene expression z scores for these genes over time. Peak IL1B expression occurs at 30min-1hr while NFKBIA is highly expressed at 1hr. ICAM1 is moderately expressed at 1hr and is highly expressed at 24hr and IFIT3 is highly expressed 24hr after stimulation with LPS.

Supp. Fig. 1. Panel A shows nucleosome distribution over a 4kb window centered over the on pol II transcription start sites (TSSs) at different time points (0hr, 5m, 30m, 1hr, 4hr, 12hr, 24hr, 48hr) following LPS stimulation. There is a 200 fold increase in signal over the central 2kb region compared to the outer edges indicating successful enrichment of promoter captured regions. Panel B displays a heatmap of Pearson's correlation coefficients for nucleosome signal in 10bp bins over promoters across the same time points. Each square in the heatmap contains the correlation value, with a color gradient indicating the strength of the correlation. Every sample has a Pearson's Correlation Coefficient above 0.87 indicating that samples are highly similar to each other.

Supp. Fig. 2. Panel A shows gene ontology (GO) terms for genes with altered nucleosome positioning at various time points (5 min, 30 min, 1 hr, 4 hr, 12 hr, 24 hr, and 48 hr) following LPS stimulation. Adjusted P values are shown beside the ontology terms, with significant enrichments represented by blue bars. The x-axis shows the -log10(p-value), representing the significance of the enrichment for each GO term, while the y-axis lists the specific GO terms along with their adjusted p-values. Panel B shows the GO terms for genes with altered nucleosome fuzziness at the same time points (5 min, 30 min, 1 hr, 4 hr, 12 hr, 24 hr, and 48 hr) following LPS stimulation. Adjusted P values are also displayed next to the ontology terms, with significant enrichments highlighted in blue bars. The x-axis shows the -log10(p-value), representing the significance of the enrichment for each GO term, while the y-axis lists the specific GO terms along with their adjusted p-values.

Supp. Fig. 3.  Panel A shows gene ontology (GO) terms for genes with increased nucleosome occupancy at various time points (5 min, 30 min, 1 hr, 4 hr, 12 hr, 24 hr, and 48 hr) following LPS stimulation. Adjusted P values are shown beside the ontology terms, with significant enrichments represented by blue bars. Panel B shows the GO terms for genes with decreased nucleosome occupancy at the same time points (5 min, 30 min, 1 hr, 4 hr, 12 hr, 24 hr, and 48 hr) following LPS stimulation. Adjusted P values are also displayed next to the ontology terms, with significant enrichments highlighted in blue bars.

Supp. Fig. 4. Panel A presents k-means clustered line plots of gene expression (Log2(FPKM+1)) for promoters with repositioned nucleosomes at various time points (0hr, 30min, 1hr, 4hr, 12hr, 24hr) following LPS stimulation. Each subplot represents different clusters of genes, with the y-axis showing gene expression levels and the x-axis showing time points. Different colored lines indicate individual gene expression trajectories within each cluster. Panel B shows k-means clustered line plots of gene expression (Log2(FPKM+1)) for promoters with altered fuzziness at the same time points following LPS stimulation. Similar to section A, each subplot represents different clusters of genes, with the y-axis representing gene expression levels and the x-axis representing time points. Various colored lines within each subplot illustrate individual gene expression trajectories within that cluster.

Supp. Fig. 5. Panel A presents k-means clustered line plots of gene expression (Log2(FPKM+1)) for promoters with increased nucleosome occupancy patterns at various time points (0hr, 30min, 1hr, 4hr, 12hr, 24hr) following LPS stimulation. Each subplot represents different clusters of genes, with the y-axis showing gene expression levels and the x-axis showing time points. Different colored lines indicate individual gene expression trajectories within each cluster. Panel B shows k-means clustered line plots of gene expression (Log2(FPKM+1)) for promoters with decreased nucleosome occupancy patterns at the same time points following LPS stimulation. Similar to section A, each subplot represents different clusters of genes, with the y-axis representing gene expression levels and the x-axis representing time points. Various colored lines within each subplot illustrate individual gene expression trajectories within that cluster.

Supp. Fig. 6. Panel A shows a bar chart displaying the Biological Process gene ontology (GO) terms with adjusted p values for common genes in the positioned sensitive clusters at 24hr and 48hr. The x-axis shows the -log10(p-value), representing the significance of the enrichment for each GO term, while the y-axis lists the specific GO terms along with their adjusted p-values. Panel B shows a bar chart of Biological Process GO terms with adjusted p values for genes in the positioned sensitivity clusters that were unique to 24hr. Similar to section A, the x-axis shows the -log10(p-value) and the y-axis lists the specific GO terms along with their adjusted p-values. Panel C shows a bar chart showing the Biological Process GO terms with adjusted p values for genes in the positioned sensitivity clusters that were unique to 48hr. The x-axis displays the -log10(p-value), and the y-axis lists the specific GO terms along with their adjusted p-values.

Supp. Fig. 7. Panels A-F show fragmaps representing the distribution of paired-end fragments from light digests by size over a 2kb region centered on the Transcription Start Site (TSS) of promoters which are separated into groups by sensitivity clustering at 0hr, 30min, 1hr, 4hr, 24hr, and 48hr following LPS stimulation. Each panel contains seven clusters (cluster1 to cluster7) with the paired-end read size (bp) on the y-axis and the TSS on the x-axis. Panel G shows bar graphs showing average nucleosome occupancy, as measured by fuzziness scores, with error bars showing 95% confidence intervals for sensitivity clusters at all time points (0hr, 5min, 30min, 1hr, 4hr, 12hr, 24hr, 48hr). Panel H represents bar graphs showing average nucleosome positioning scores, as measured by summit scores, for sensitivity clusters at all time points (0hr, 5min, 30min, 1hr, 4hr, 12hr, 24hr, 48hr) with error bars representing 95% confidence intervals.

Supp. Table 1
List of genes with one or more nucleosomes within promoter regions with altered positioning at one or more time points following LPS stimulation compared to unstimulated cells.  We defined nucleosomes as being repositioned if there was a shift in the nucleosome summit position greater than 80bp between time points. Each column in the table includes a list of genes for a given time point which is listed in the first row of the column. There is no 0hr time point listed as that was the baseline used to compare with all other time points.

Supp. Table 2
List of genes with one or more nucleosomes within promoter regions with altered fuzziness at one or more time points following LPS stimulation compared to unstimulated cells. We considered a change in nucleosome fuzziness to occur if there was a log2 fold change greater than 1.5 between conditions combined with a p-value and FDR under 0.05. Each column in the table includes a list of genes for a given time point which is listed in the first row of the column. There is no 0hr time point listed as that was the baseline used to compare with all other time points.

Supp. Table 3
List of genes with one or more nucleosomes within promoter regions with increased occupancy at one or more time points following LPS stimulation compared to unstimulated cells. We defined alterations in nucleosome occupancy as having a log2 fold change greater than 5 with a p-value and FDR under 0.05 for any time point compared to the basal occupancy seen in 0hr. Each column in the table includes a list of genes for a given time point which is listed in the first row of the column. There is no 0hr time point listed as that was the baseline used to compare with all other time points.

Supp. Table 4
List of genes with one or more nucleosomes within promoter regions with decreased occupancy at one or more time points following LPS stimulation compared to unstimulated cells. We defined alterations in nucleosome occupancy as having a log2 fold change less than 5 with a p-value and FDR under 0.05 for any time point compared to the basal occupancy seen in 0hr. Each column in the table includes a list of genes for a given time point which is listed in the first

row of the column. There is no 0hr time point listed as that was the baseline used to compare with all other time points.
